# Systematic proteomics identifies a conserved mechanism for polar targeting of plant cortical proteins

**DOI:** 10.64898/2026.08.17.745269

**Authors:** Evgeniya M. Pukhovaya, Catherine Albrecht, Maritza van Dop, Victor Jones, Mark Roosjen, Andriy Volkov, João Jacob Ramalho, Sumanth Mutte, Chang Su, Helen Strutt, Joyce C.M. Meiring, Anna Akhmanova, David Strutt, Dolf Weijers

## Abstract

Multicellular development is tightly coupled to the polarization of individual cells, which partitions polar proteins along the cell cortex and can control asymmetric cell division, anisotropic growth, local differentiation or physiology. Mechanisms driving cell polarization have been described in fungi and animals, but these lack counterparts in plants. While several polarized proteins have been identified in plants, the overall mechanisms guiding their polar localization are poorly characterized. Through iterative affinity proteomics on the recently identified SOSEKI polar proteins, we discovered a network of polar proteins that is conserved in the flowering plant Arabidopsis and the liverwort Marchantia. We next used this collection of novel polarized proteins for systematic proximity ligation proteomics in these two species to map their global polar proteome. We identified a subfamily of polarized Protein S-acyl transferase (PAT) enzymes that are required for membrane targeting of Arabidopsis SOSEKI proteins. Using human and fruit fly models, we showed that PAT19 is sufficient for membrane targeting of SOSEKI1, likely through direct palmitoylation. This work demonstrates a conserved mechanism for polar protein targeting in plants and offers a resource for studying polar protein localization.

## Introduction

Cell polarity is key to generating asymmetrical spatial cell organization, including the polar localization of organelles and specific molecules – proteins, lipids and nucleic acids^1–3^. In multicellular organisms, cell polarity can instruct the orientation of cell division and cell growth, and allows for directional transport and subcellular functional specialization^4–7^. In a multicellular context, the coordinated polarization of cells is key to body axis formation^8,9^. Cells in all domains of life show features of polarization^10–13^, yet the molecules that are polarized and the mechanisms that guide this process are very different. As multicellularity evolved independently in animals, plants, fungi, and red and brown algae^14^, so did the integration of cell polarity in their developmental programs. Thus, fully understanding principles of cell polarity requires the study of multiple domains of life. Land plants have highly complex body architectures with many cells and cell types that are highly polarized. Most plant cells in vegetative tissues have a polyhedral shape with distinct faces. Such cell faces are often defined along organ or organismal axes, and therefore reflect apical and basal (following organ elongation direction), inner and outer, or lateral (oriented towards the organ surface), and radial (facing similar cells)^12^. Polar proteins can be concentrated at the plasma membrane (PM) at these faces, generating polar PM domains^15,16^. The study of plant cell polarity has been focused extensively on the polarly localized PIN-FORMED auxin efflux transporters (PINs)^17^, and this has identified steps in vesicle trafficking that mediate polar targeting of these transmembrane proteins18. Importantly, PIN protein polarity is the net result of dynamic cycling between the PINs and intracellular compartments. This process is controlled by vesicle recycling (through GNOM ARF-GEF^18^), protein degradation (through WAVE3^19^), and restricting PIN lateral diffusion (through MEL/MAB^20^). This is a highly dynamic process that is affected by various external signals such as light (through NPH3^21^) and gravitropic changes (through RLD and LAZY^22,23^). Furthermore, cell polarity in plants has been connected to vascular cell differentiation (OPS^24^, BRX^25^) and asymmetric division in the stomatal lineage (BASL^5^, BRXf^26^, POLAR^27^). All these processes and polar proteins are characterized by highly dynamic changes in protein localization. Hence, a major outstanding question in plant cell polarity is what stable cellular polarity landmarks offer reference to the polarity system, and how such stable patterns are generated.

SOSEKI (SOK) polar proteins localize to specific cell edges^28^, thus occupying apical-basal and lateral polar PM domains. In contrast to most other known plant polarity proteins, the patterns of SOK localization are highly stable in interphase cells^28^, and is also evident upon ectopic expression. SOK proteins are deeply conserved in land plants^29^ and contain a DIX oligomerization domain that is structurally and functionally homologous to the DIX domain in metazoan Dishevelled protein^29^. In analogy with Dishevelled, SOK may act as a polar, polymeric scaffold that recruits other proteins, such as ANGUSTIFOLIA (AN), to polar cortical domains^29,30^. Given the unique localization of SOK proteins at the intersection of apical-basal and lateral cell axes, we hypothesize that SOK proteins connect to the components of various polar PM domains in plant cells. Dissecting their targeting mechanism may therefore reveal novel regulators of polar protein targeting in plants.

Here, we use SOK proteins as a starting point for proteomic dissection of their interactions. There is very little proteomic information on plant cell polarity, but those that were reported suggest connections between polar proteins. For example, SOK3 and AN were found in an OPL proximity labeling interactome in stomata^31^, and split-TurboID on BRXL2 and its interactor BASL^32^ identified new components of basal polarity in the hypocotyl and root and connected them to the response to light. Thus, proximity localization proteomics can reveal relevant interactions among polar proteins and help connect these to other cellular processes.

Here, we generate an iterative interactome of the SOK protein network. We use a set of newly identified polar proteins to perform a systematic proximity ligation proteome survey of the polar proteome. By generating datasets in the flowering plant Arabidopsis and the liverwort Marchantia, we discover a conserved set of polar proteins and identify a novel protein palmitoylation-based targeting module. We find this module to be required for SOK targeting in Arabidopsis and sufficient for membrane targeting in heterologous animal systems. Thus, through systematic proteomics, our work identifies a mechanism for the membrane targeting of cortical polar proteins and offers a framework for studying plant cell polarity.

## Results

### Identification of a SOSEKI-centered protein network

We previously performed immunoprecipitation followed by mass spectrometry (IP-MS) on YFP-tagged Arabidopsis SOK1, 2 and 3 proteins, which revealed that SOK proteins can recruit each other and the ANGUSTIFOLIA (AN) effector in a polar manner^29^. To further explore the protein context of SOK polarity, we re-analyzed these IP-MS datasets (**Fig. 1A-C**). This analysis recovered a set of proteins associated with one or more SOK proteins (**Supplementary data 1**). We next generated YFP fusions for two interactors: SOSEKI1-INTERACTING BTB1 / NPH3/RPT2-LIKE2 (SIB1/NRL2) and DYNEIN LIGHT CHAIN1 (DLC1) – which were selected based on their identification in multiple of the above datasets. IP-MS on these lines and on AN-GFP recovered an overlapping set of proteins as for SOK experiments. These included SOK, AN and DUAL-SPECIFICITY TYROSINE PHOSPHORYLATION-REGULATED KINASES (DYRKP1 and DYRKP2A/B). Additionally, we identified two SIB1 homologs (SIB2/NRL16 and SIB3/NRL29), a putative cytosolic kinase (SIB1-INTERACTING CYTOSOLIC KINASE; SICK), not homologous to any previously described kinase families, a receptor-like kinase (SIB1-INTERACTING TRANSMEMBRANE KINASE1; SITK1) and the γ-tubulin ring complex adaptor NEDD1 (**Fig. 1D-F**). We next generated an anti-YFP IP-MS dataset for SICK-TurboID-YFP (**Fig. 1G**). This recovered SIB1, SIB2 and SIB3 proteins, as well as a SITK1 homolog (SITK2).

**Fig. 1.**
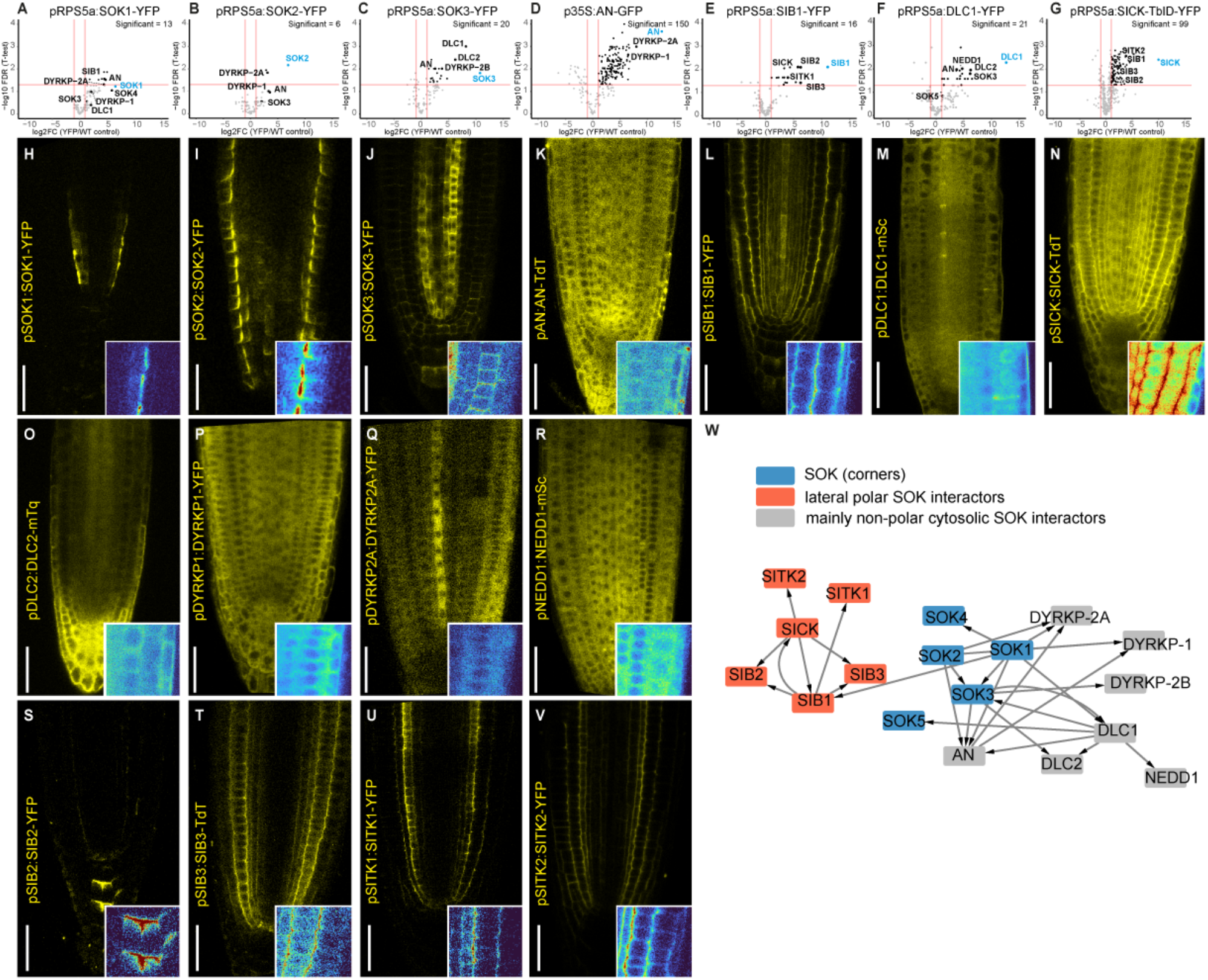
Iterative proteomics identifies a SOK-centered polar protein network. (**A**-**G**) Volcano plots depicting significantly (Benjamini-Hochberg false discovery rate (FDR) < 0.05, fold change (logFC) > 1) enriched proteins in IP-MS experiments comparing technical triplicates of 7-days-old seedling roots expressing (**A**) pRPS5a:SOK1-YFP, (**B**) pRPS5a:SOK2-YFP, (**C**) pRPS5a:SOK3-YFP, (**D**) p35S:AN-GFP, (**E**) pRPS5a:SIB1-YFP, (**F**) pRPS5a:DLC1-YFP, (**G**) pRPS5a:SICK-TurboID-YFP against WT Col-0 roots. The overexpressed transgenic protein is highlighted in blue, the proteins of interest identified in multiple datasets are labeled in black. The number of significantly enriched interactors above the threshold (n) in each line is indicated. (**H**-**V**) Confocal images of roots of 7-days-old Arabidopsis seedlings expressing (**H**) pSOK1:SOK1-YFP, (**I**) pSOK2:SOK2-YFP, (**J**) pSOK3:SOK3-YFP, (**K**) pAN:AN-TdTomato, (**L**) pSIB1:SIB1-YFP, (**M**) pDLC1:DLC1-mScarlet, (**N**) pSICK:SICK-TdTomato, (**O**) pDLC2:DLC2-mTurquoise, (**P**) pDYRKP1:DYRKP1-YFP, (**Q**) pDYRKP2A:DYRKP2A-YFP, (**R**) pNEDD1:NEDD1-mScarlet, (**S**) pSIB2:SIB2-YFP, (**T**) pSIB3:SIB3-TdTomato, (**U**) pSITK1:SITK1-YFP and (**V**) pSITK2:SITK2-YFP. Scale bars: 40 µm. Bottom right corner of each image shows an inset of a few epidermal and cortex cells (with the exception of pericycle for SOK1 and columella for SIB2) in false color intensity scale. (**W**) Protein network summarizing the IP-MS interactions identified in (**A**-**G**). Arrows point towards the proteins found as immunoprecipitated interactors in the corresponding transgenic lines. The SOK family is highlighted in blue, laterally polar interactors in red, and non-polar cytosolic interactors in gray.

SOK proteins predominantly localize to cell edges, while AN localizes to the cytosol in a non-polar manner (consistent with the previous research, when SOK was not overexpressed^28,29^) (**Fig. 1H-K**). To check the localization of the newly identified interactors, we analyzed fluorescent fusion proteins in roots. We found SIB1 and DLC1 to show polar localization (**Fig. 1L-M**), with SIB1 being laterally polarized in the ground tissue and DLC1 being apical-basally polarized in some vascular cells. We found SICK also to be polarly localized (**Fig. 1N**), showing a lateral pattern in the ground tissues similar to SIB1; SICK additionally had a high fraction of cytosolic non-polar protein. In contrast, DLC2, the DYRKP kinases, and NEDD1 showed only cytosolic localization (**Fig. 1O-R**). SIB2, SIB3, SITK1 and SITK2 were polarly localized (**Fig. 1S-V**); SIB2 was apical in root cap cells, while SIB3, SITK1 and SITK2 showed a lateral pattern in the ground tissues similar to SIB1. We summarized the IP-MS identifications of all proteins described above in a network that represents the reproducibly identified interactors for each bait (**Fig. 1W**).

The co-IP and co-localization data for SIB1-3, SICK, and SITK1-2 indicate that these proteins form a tightly connected lateral polar complex. The SOK interactome, by contrast, showed no co-localization of the identified proteins with SOK in cell corners. Instead, SOK associated with two distinct protein pools - one at the PM on the lateral side of the cell, and one in the cytosol - suggesting that a cytosolic and a PM-associated fraction of SOK engage different partners. This set of interactors likely includes regulators of SOK localization as well as proteins linking SOK to their cellular functions.

### Mapping of the polarized proteome through systematic proximity ligation

With the identification of a set of novel polarized proteins in Arabidopsis roots (**Fig. 1**), we aimed to generate a resource for the systematic identification of polarized proteins. We employed a proximity ligation strategy (**Fig. 2A**) to develop an atlas of association with a range of polarized proteins. We fused the TurboID biotin ligase enzyme^33^ and YFP (TbID-YFP) to a large range of proteins with different subcellular polar localizations (**Fig. 2B**). We included several SOSEKI-network proteins: SOK1, SOK2, SOK3, SOK4, SOK5, AN, DLC1, SIB1, SICK, SITK1. In addition, we included previously reported polar proteins that bear no known relation to SOK: NIP5;1, BOR1, BRX, POLAR, OPS, PIN2, BASL (**Fig. 2B**). This set included proteins of various polarity classes, likely representing different targeting mechanisms: those that have dual cytosolic and polar membrane localization (AN, DLC1, SICK), cortical polar membrane proteins (SOK, SIB1, BRX, POLAR, OPS, BASL) and polar transmembrane proteins (SITK1, NIP5, BOR1, PIN2). For proximity labeling, we generated transgenic Arabidopsis lines expressing each protein-TbID-YFP fusion from the meristematic pRPS5A promoter^34^ (**Fig. 2C-Q**). For a control for non-specific “sticky” proteins and highly abundant PM proteins, we also transformed an empty vector pRPS5a:TbID-YFP without the bait protein (later referred to as a free ligase control) (**Fig. 2C**) and plasmids carrying two non-polar transmembrane PM proteins LTI6B and PIP2A (**Fig. 2P,Q**). We could not recover stable pRPS5A-driven Arabidopsis lines carrying PIN2, BOR1, SOK2, OPS or BASL, likely due to lethal phenotypes caused by misexpression or transgene silencing. For SOK1, SOK4 and AN, we also observed a significant reduc tion of expression in subsequent generations, likely due to transgene silencing. In all stable lines, TbID protein fusions localized as predicted in the root meristem (**Fig. 2C-Q**). Over- and misexpression rendered some polar proteins, such as NIP5;1 (**Fig. 2O**), less prominently polar. In contrast, DLC1 (**Fig. 2I**) was seen in the apical/basal side of the PM more often than in native promoter expression lines.

**Fig. 2.**
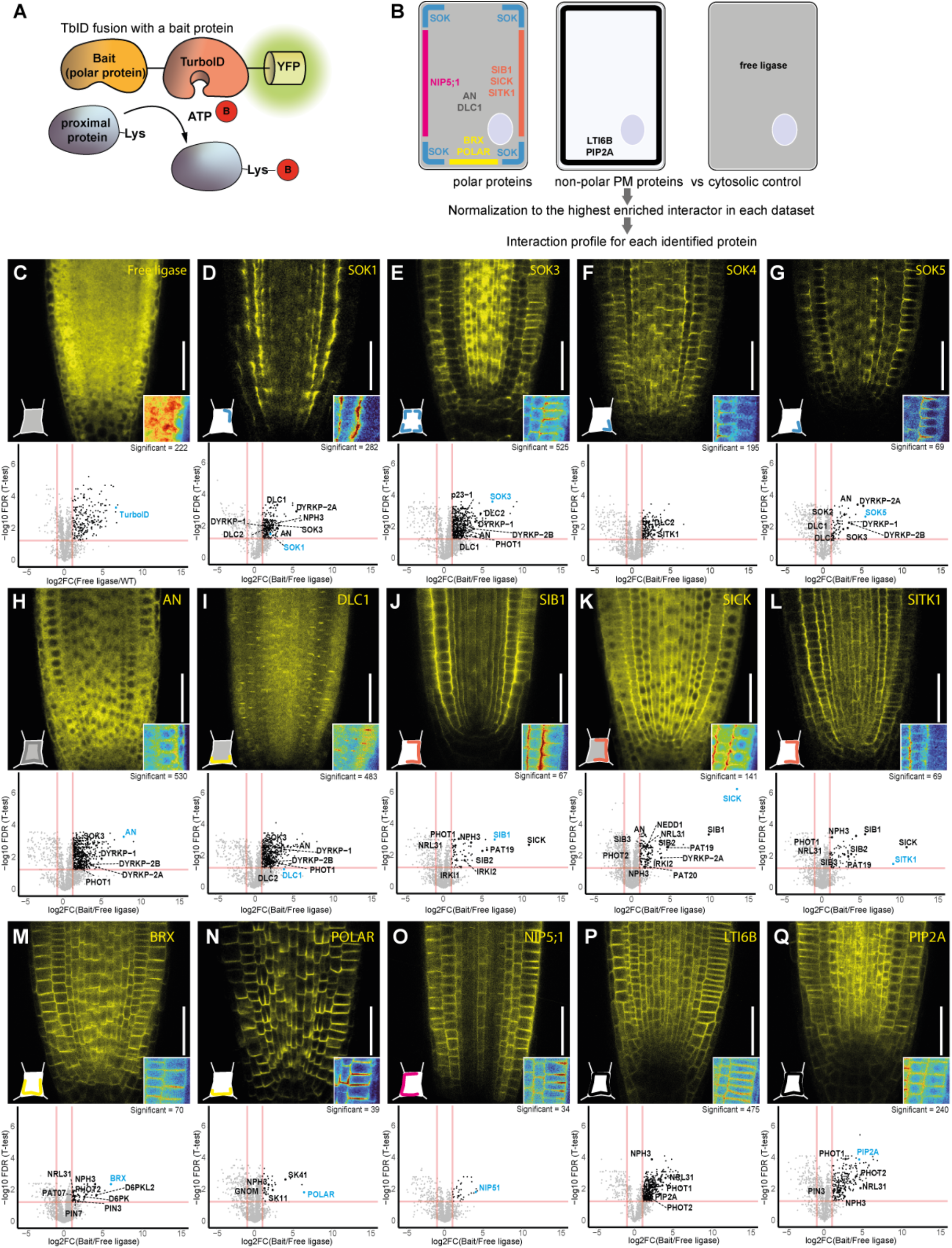
Generation of a comprehensive proximity labeling dataset for Arabidopsis polar proteins. (**A**) Schematic representation of a transgenic fusion between TurboID (TbID) and a protein of interest (bait) and the labeling reaction. (**B**) Schematic representation of Arabidopsis root cells expressing TbID-YFP targeted to various cellular polar domains via the fusion to known polar proteins. Protein name colors represent distinct polar domains. After the datasets from each transgenic line were generated, the enrichment against non-polar free ligase control was calculated, and the logFC enrichment values were normalized to the highest logFC in each dataset. The combination of normalized logFC values for each identified interactor in each dataset was called an interaction profile. (**C**-**Q**) Confocal images of 7-days-old roots expressing pRPS5a:bait-TbID-YFP (scale bar 40 µm) and the volcano plots of corresponding proximity labeling MS data for (**C**) pRPS5a:TbID-YFP, (**D**) pRPS5a:SOK1-TbID-YFP, (**E**) pRPS5a:SOK3-TbID-YFP, (**F**) pRPS5a:SOK4-TbID-YFP, (**G**) pRPS5a:SOK5-TbID-YFP, (**H**) pRPS5a:AN-TbID-YFP, (**I**) pRPS5a:DLC1-TbID-YFP, pRPS5a:SIB1-TbID-YFP, (**K**) pRPS5a:SICK-TbID-YFP, (**L**) pRPS5a:SITK1-TbID-YFP, (**M**) pRPS5a:BRX-TbID-YFP, (**N**) pRPS5a:POLAR-TbID-YFP, (**O**) pRPS5a:NIP5;1-TbID-YFP, (**P**) pRPS5a:LTI6B-TbID-YFP and (**Q**) pRPS5a:PIP2A-TbID-YFP. Volcano plots depict the number of significantly (Benjamini-Hochberg false discovery rate (FDR) < 0.05, fold change (logFC) > 1) enriched proteins identified between technical triplicates. Bait protein is highlighted in blue. The names of proteins with previously shown physical or functional connection to the bait, as well as several interactors discussed in the next sections, are labeled in black. Bottom right corner of each image shows in inset of a few epidermal and cortex cells from the right side of the root with false-color intensities. Bottom left corner shows schematic representation of a root cell from the left side, depicting the major localization of each polar protein, color code from panel (**A**).

Proximity ligation requires addition of external biotin for efficient labeling^35^. We treated the SOK5, SIB1, NIP5;1 and LTI6B-TbID-YFP lines with 50 µM biotin and found that (polar) membrane localization was largely retained after 6 and 24 hours of treatment (**Fig. S1A-P**), even though SOK5 and NIP5;1 localization seemed to be more sensitive to flooding, and thus the treatment might affect the native protein environment. We next performed mass spectrometry on the SIB1-TbID-YFP line after 6h (**Fig. S1Q**) and 24 h (**Fig. S1R**) and compared the results (**Fig. S1S**). After 24h, we identified 146 more interactors compared to the 6h treatment, including DLC1 (secondary interactor not found in IP-MS of SIB1 but found in SOK IP-MS), and therefore chose the 24h time point for the systematic proximity ligation on all proteins.

We next performed systematic proximity ligation on all 15 baits, including the free ligase control (pRPS5a:TbID-YFP) and the two non-polar plasma membrane controls (pRPS5a:LTI6B/PIP2A-TbID-YFP). We performed biological (independent experiments) for 7 baits and technical (separate samples within a single experiment) replications for all baits (**Supplementary data 2**). The free ligase control recovered many significantly enriched proteins compared to non-transgenic plants (**Fig. 2C**), representing the background labeling. Thus, free ligase was chosen as a reference to quantify the fold change (logFC) of a bait of interest. In this systematic analysis, we recovered most of the previously described (**Fig. 1W**) SOK network components in the corresponding bait transgenic lines (**Fig. 2D-L**). While the order of identification was not the same as in the IP-MS, it highlighted the higher radius of action and the advantage of combining both techniques. Likewise, for other polar proteins, we identified known interactors (e.g. D6PK and PIN with BRX^25,36^; **Fig. 2M**) or proteins that could be potentially involved in their function (Shaggy-like kinases SK11 and SK41 with POLAR^27^, **Fig. 2N**) or polar targeting (GNOM with PO-LAR, **Fig. 2N**). Among the different baits and biological replicates for the same bait, we observed substantial variability in the number of significantly enriched proteins (**Fig. 2C-Q**). This can be explained by the difference across biological replicates, as well as by the data-dependent MS acquisition mode that selects the most abundant peptides for MS2 fragmentation in each experiment independently. We also observed that logFC distributions differed between datasets, likely due to variations in bait expression between independent transgenic lines. To account for the differences between datasets, we normalized values by dividing each logFC by the maximum logFC in the relevant dataset (**Fig. 2A**). We referred to the set of normalized logFC values from each dataset for each identified protein as an interaction profile of this protein. Interaction profiles allowed us a cross-dataset comparison while preserving relative information about protein enrichment. Joining all datasets identified 2511 (logFC > 0, FDR < 0.05) or 2184 (logFC > 1, FDR < 0.05) significantly enriched proteins in at least one of the proximity labeling experiments, most of which were identified in the cell polarity network for the first time. The data can be downloaded or browsed in a Shiny app (https://weijerslab.shinyapps.io/polar_protein_proxitome/).

### Identification of a cluster of novel polarized proteins

In contrast to conventional IP-MS or proximity labeling experiments that use one or a few bait proteins, we leveraged the power of the large set of baits with distinct, yet partially overlapping localizations. We reasoned that novel polarized proteins could be identified through global patterns of association with the entire set of baits. For example, a SOK-specific partner protein would be expected to be enriched in most or all of the SOK datasets. To reduce the complexity of the dataset and identify dominant patterns, we used k-means clustering for all the 27 datasets based on the normalized logFC value of each significantly enriched proximity interactor (**Fig. 3A**). The optimal number of clusters (n=20) was determined using the Elbow method (**Fig. S2A**). Among the 20 clusters, cluster 4 was remarkable since its members were enriched in SOK1, SOK3, SOK5, AN, DLC1 and SICK samples (**Fig. 3B**). Since all these baits were derived from SOK IP-MS experiments (**Fig. 1**), we considered this a “SOK cluster”. Apart from the bait proteins themselves, this cluster included the DYRKP kinases, which were previously identified only as interacting with SOK and AN. We consider the identification of this cluster a validation of our approach, and we take this as evidence that proximity labeling allows identification of indirect or transient interactions. In the clustering, cluster 8 was specifically enriched in laterally localized SOK complex components SIB1, SICK and SITK (**Fig. 3B**). Apart from baits and their previously identified interactors SIB2 and SIB3, it included 2 other proteins - NPH3 (NON-PHOTOTROPIC HYPOCOTYL3) and PAT19 (S-PROTEIN-ACYL TRANSFERASE 19). We next explored the robustness of cluster identification by using different statistical thresholds and clustering parameters (**Shiny app Clustering Tab**). This lateral cluster remained stable using various clustering parameters: SIB1-2, SICK, PAT19 and NPH3 remained clustered together (**Fig. S2B**). With some parameters, the IAP1 (IMMUNE-ASSOCIATED NUCLEO-TIDE-BINDING9 ASSOCIATED PROTEIN1) also clustered together with the lateral module (**Fig. S2B**).

**Fig. 3.**
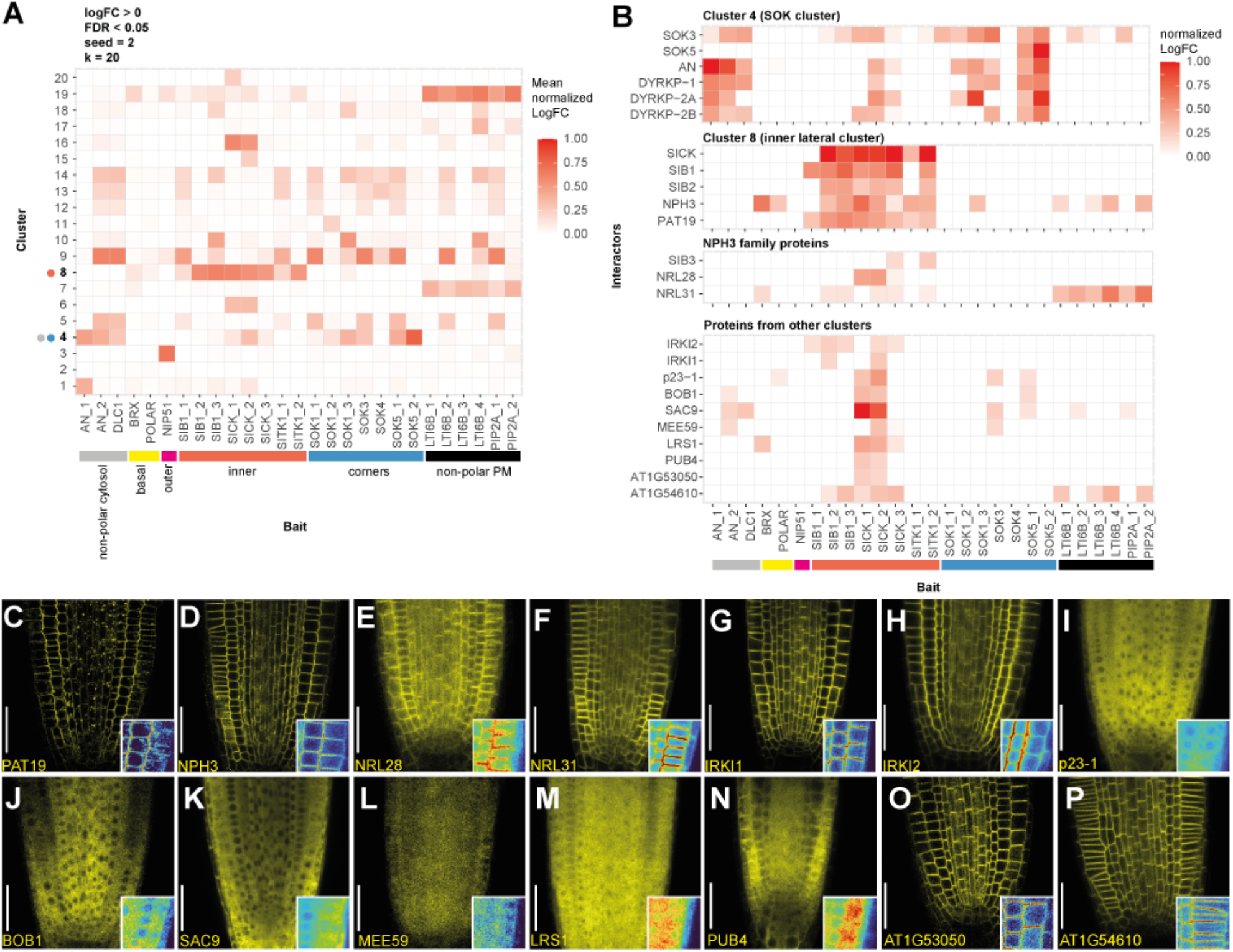
Proximity labeling-based proteomics identifies new polar proteins in roots. (**A**) k-means clustering of the interaction profiles of significantly enriched interactors identified in the polar proteins proximity labeling dataset. Mean normalized logFC across all the proteins in each cluster is depicted in the heatmap. Horizontal bar below the heatmaps shows the localization of the bait proteins in root cells. Circles next to clusters 4 and 8 represent the localization of bait proteins where these clusters are mostly enriched. (**B**) Detailed heatmaps of clusters and groups of proteins of interest depicting normalized logFC values (interaction profiles) for each protein: cluster 4 and cluster 8 from panel (**A**), NPH3 family proteins which were not part of cluster 8, and all the protein candidates selected outside clusters 4 and 8. (**C**-**P**) Confocal images of 7-day-old roots expressing selected candidate proteins (scale bar 40 µm). (**C**) pRPS5a:PAT19-TbID-YFP (**D**) pRPS5a:NPH3-TbID-YFP (**E**) pRPS5a:NRL28-TbID-YFP (**F**) pRPS5a:NRL31-TbID-YFP (**G**) pRPS5a:IRKI1-TbID-YFP (**H**) pRPS5a:IRKI2-TbID-YFP (**I**) pRPS5a:p23-1-TbID-YFP (**J**) pRPS5a:BOB1-TbID-YFP sac9 pSAC9:mCitrin-SAC9 (**L**) pRPS5a:MEE59-TbID-YFP (**M**) pRPS5a:LRS1-TbID-YFP (**N**) pRPS5a:PUB4-TbID-YFP (**O**) pRPS5a:AT1G53050-TbID-YFP (**P**) pRPS5a:AT1G54610-TbID-YFP. Bottom right corner of each image shows an inset of a few epidermal and cortex cells from the right side of the root with false-color intensities.

Since IAP1 was reported as a cytosolic regulator of immune response^37^, we considered it unlikely to be directly relevant to cell polarity. The other novel cluster components, PAT19 and NPH3, had by contrast already been assigned functions that could connect them to polar proteins: NPH3 domain-containing proteins might control cellular trafficking^38^, while PATs catalyse membrane targeting of their substrates and could therefore participate in the polar targeting of pro teins^39^. We studied the localization of PAT19 and NPH3 in roots (**Fig. 3C-D**). While localizing to the PM and to the endomembrane compartments, PAT19 was polarized towards the inner lateral side of epidermal and cortex cells (**Fig. 3C**), copying the polarity of SIB1 and SICK. In contrast, NPH3 was not polarized in root cells (**Fig. 3D**). As NPH3 belongs to the same protein family as SIB1/2/3, we localized other members of this NRL family found in our datasets, but we found these – NRL28 and NRL31/NCH1^40,41^ – to also be nonpolar (**Fig. 3E-F**). Thus, only the members of the SIB clade were polar among the NPH3 family proteins identified in our proximity labeling screen.

In the remaining clusters, interaction profiles did not allow us to make functional hypotheses. Yet, these clusters contained polar proteins whose hypothetical functions or high enrichment level made them promising for further functional analysis. We selected 10 proteins enriched in more than 1 dataset and studied their localization in roots (**Fig. 3B, G-P**). IRKI1 (AT5G12900, IRK-interacting protein 1) was previously shown to interact directly with polar kinase IRK^42^. Indeed, we identified IRKI1 in the SIB1 and SICK datasets, which also interact with SITK1/2 – homologs of IRK^43^. While IRKI1 was not polar when expressed from the RPS5A promoter (**Fig. 3G**), its close homolog IRKI2 (AT1G12330) localized to the lateral polar membrane (**Fig. 3H**), resembling SIB1, SICK, SITK, and PAT19. Proteins p23-1, BOB1, SAC9 and MEE59 were previously related to other types of polarity: apical/basal (p23^44^, BOB1^45^), directional vesicle trafficking during cell division (SAC9^39^), or found to be indispensable for early development (BOB1^45^, MEE5946). Our transgenic lines for these constructs did not demonstrate any polar signal (**Fig. 3I-L**). Additionally, we studied the localization of 4 proteins highly enriched in the laterally localized SICK kinase datasets – LATERAL ROOT STIMULATOR LRS1, plant U-box 4 PUB4, and 2 putative protein kinases AT1G53050 and AT1G54610 (**Fig. 3M-P**). All of them appeared to be non-polar. In summary, initial analysis of our systematic proximity labeling approach identified 2 new protein families with laterally polarized members – PAT19 and IRKI.

### Deep conservation of a polarized protein network

SOK proteins are deeply conserved across land plants and are polarized in all studied cases, including in the liverwort *Marchantia polymorpha* and the moss P*hyscomitrium patens*^*2*9^. In Marchantia gemmae, SOK points towards the apical notch, resembling the corner localization of Arabidopsis SOK1 in roots (**Fig. 4A**). This suggests that not only SOK proteins but also their polar targeting mechanisms likely emerged early during land plant evolution. Very few polarized proteins have been isolated in bryophytes, which complicates comparisons across land plants. We searched for homologs of the core Arabidopsis SOK-associated protein network (**Fig. 1W**), identified single homologs for SIB/NPT and SICK in Marchantia and found these to be polarized relative to the apical notch in gemmae, when expressed as TbID-YFP fusions (**Fig. 4A-C**). Note that MpSIB and MpSICK show similar localization, yet opposite to MpSOK (**Fig. 4A-C**).

**Fig. 4.**
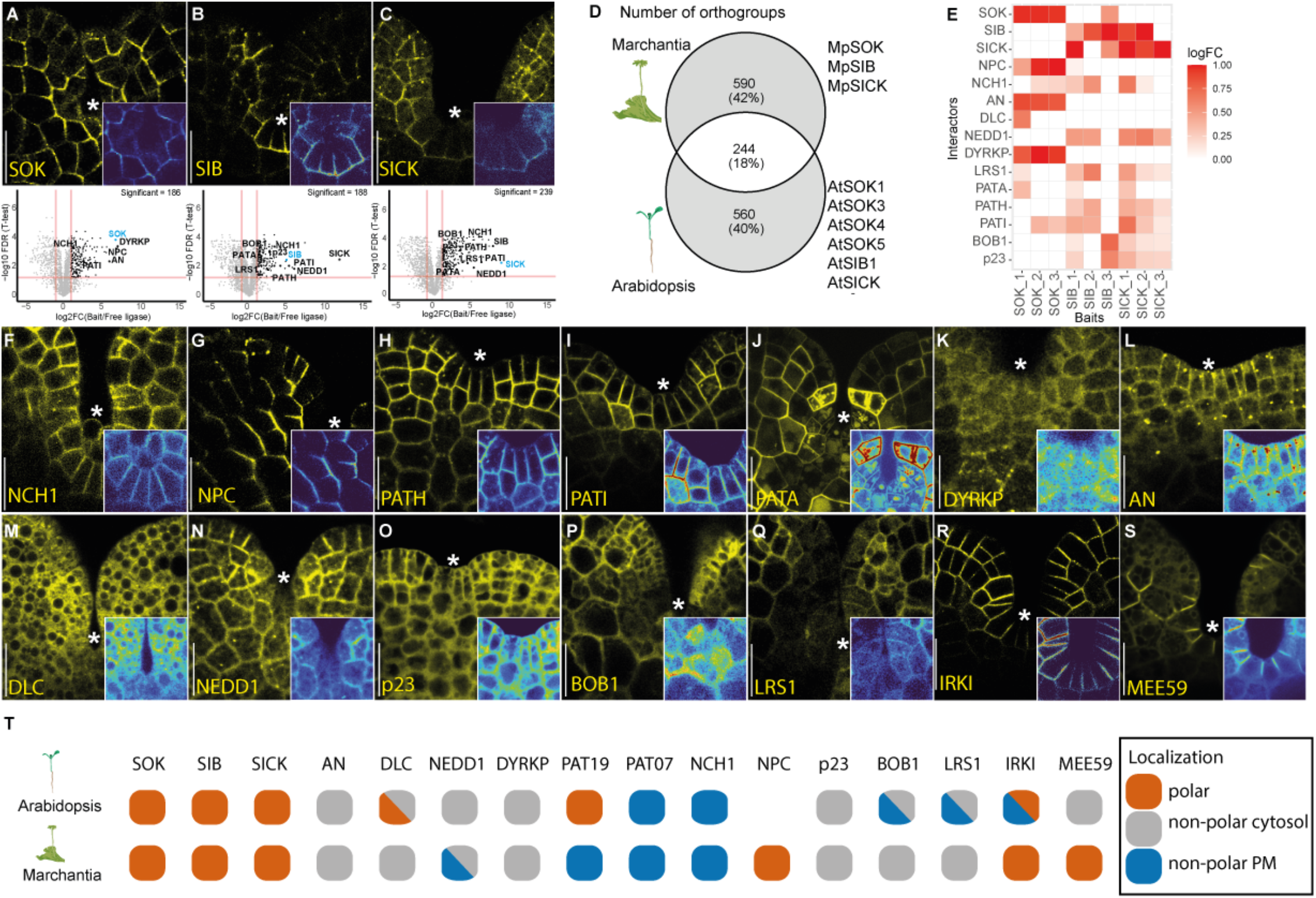
Proximity labeling in Marchantia reveals homologous polarity networks. (**A**-**C**) Confocal images of Marchantia gemmae expressing pEF1:bait-TbID-YFP (scale bar 20 µm) and volcano plots of corresponding proximity labeling MS data for (**A**) pEF1:SOK-TbID-YFP, (**B**) pEF1:SIB-TbID-YFP and (**C**) pEF1:TbID-YFP-SICK. Volcano plots depict the number of significantly (Benjamini-Hochberg false discovery rate (FDR) < 0.05, fold change (logFC) > 1) enriched proteins identified between technical triplicates. Bait protein is highlighted in blue. Bottom right corner of each image shows an inset of a few apical notch cells with false color intensities. Asterisk indicates apical notch. (**D**) Venn diagram showing intersections of orthologous groups of proteins (orthogroups) statistically enriched in Arabidopsis or Marchantia proximity labeling datasets for the protein families SOK, SIB and SICK. Proteins were filtered according to their functional annotation to exclude common mass-spec contaminants. Shared protein groups and corresponding proteins in 2 species are listed in Supplementary data 4. (**E**) Heatmap depicting normalized logFC values from Marchantia proximity labeling data for some protein candidates found in the orthogroup intersections of Arabidopsis and Marchantia datasets. (**F**-**S**) Confocal images of Marchantia gemmae expressing selected proteins from the families identified in both Marchantia and Arabidopsis proximity labeling datasets (scale bar 20 µm). (**F**) pEF1:NCH1-TbID-YFP, (**G**) pEF1:NPC-TbID-YFP, (**H**) pEF1:PATH-TbID-YFP, (**I**) pEF1:PATI-TbID-YFP, (**J**) pEF1:PATA-TbID-YFP, (**K**) pEF1:DYRKP-TbID-YFP, (**L**) pEF1:TbID-YFP-AN, (**M**) pEF1:DLC-TbID-YFP, (**N**) pEF1:NEDD-TbID-YFP, (**O**) pEF1:p23-TbID-YFP, (**P**) pEF1:BOB1-TbID-YFP, (**Q**) pEF1:LRS1-TbID-YFP, (**R**) pEF1:IRKI-TbID-YFP, (**S**) pEF1:MEE59-TbID-YFP. Bottom right corner of each image shows an inset of a few apical notch cells with false color intensities. Asterisk indicates apical notch. (T) Protein localization summary in Arabidopsis and Marchantia, based on this figure, **Fig. 1** and **Fig. 3**.

We next generated TbID fusions for MpSOK, MpSIB/NPT and MpSICK, and performed proximity labeling experiments in young thalli (**Fig. 4A-C, Supplementary data 3**). Across all replicates, we identified 1579 proteins as being enriched (logFC > 0, FDR < 0.05). To compare these proteins with the 2089 enriched proteins in proximity ligation experiments of the Arabidopsis orthologs (SOK1, SOK3, SOK4, SOK5, SIB1, SICK; **Fig. 2**), we assigned each protein to an orthogroup^47^. We additionally assigned GO terms to all the Arabidopsis proteins belonging to these orthogroups, and manually excluded proteins with generic GO terms. Out of the 1394 remaining orthogroups, 18% (244) were shared between the two species (**Fig. 4D**). Several proteins we had identified in Arabidopsis as part of the SOK network were found in the overlap (**Fig. 4E**). MpSICK was highly associated with MpSIB, and we found several other NPH3 family proteins to be enriched in Marchantia. When localized, as in Arabidopsis, the MpNCH1 protein (homolog of Arabidopsis NRL31, NRL28) was not polarized (**Fig. 4F**). We additionally found another family member, the NPC protein, to be enriched and localized polarly (**Fig. 4G**). Thus, NPH3 family members are conserved components of plant polarity systems.

Strikingly, we identified PAT19 homologs (MpPATH, MpPATI) to be enriched in the Marchantia proximity ligation data, and found these proteins to be PM-localized without obvious polarity (**Fig. 4H-I**). In addition, the Marchantia dataset also included the non-polar PM-localized MpPATA (**Fig. 4J**), while in Arabidopsis, its non-polar homolog PAT07 (**Fig. S2C-D**) was identified in the BRX dataset.

Furthermore, the Marchantia dataset also contained homologs for DYRKP, AN, DLC and NEDD1 (**Fig. 4K-N**). We generated fluorescent fusions, and found all to be localized to the cytosol, as in Arabidopsis. Thus, both cytosolic and PM-localized components of the Arabidopsis SOK network appeared to be conserved in both species. We also identified other common interactors that were previously connected to polarity processes (p23, BOB1, LRS1) (**Fig. 4O-Q**) and found these not to be polarized, as in Arabidopsis. Finally, we studied the localization of the homologs of two Arabidopsis proteins that were not identified in the Marchantia dataset: IRKI and MEE59. Both IRKI (**Fig. 4R**) and MEE59 (**Fig. 4S**) localized polarly to the sides of apical notch cells, showing a type of polarity orthogonal to the SOK complex.

In summary, we discovered a deep homology in the protein networks connected to cell polarity across land plants. In several cases, polar protein localization remain deeply conserved (**Fig. 4T**).

### Identification of a clade of polarized protein S-acyltransferases

Through systematic proteome analysis in Arabidopsis and Marchantia, we would expect to identify both regulators of polar protein targeting and effectors that connect polar complexes to cellular function. We here focus on the former and explore the function of the PAT enzymes associated with polarized proteins, given that PAT enzymes can target substrate proteins for membrane localization through the addition of a palmitoyl group to target cysteines^39^.

First, we asked if polar localization in the Arabidopsis PAT family is unique to PAT19. To this end, we constructed a phylogeny (**Fig. 5A**) and identified 8 clades that diverged early during land plant evolution. We selected Arabidopsis representatives of most of the subclades for analysis of subcellular localization by expressing all from the RPS5A promoter. Out of all the tested PATs, only the members of clade VIII (PAT19/20/21/22) showed polarized lateral localization (**Fig. 5B-I**). When expressed from their native promoters, PAT19, 20 and 22 showed overlapping polarization towards lateral cell membranes in roots, as well as varying degrees of endomembrane localization (**Fig. 5C,E,I**). No PAT21 expression could be detected in roots, confirming the recent report^48^ (**Fig. 5G**). Two Marchantia orthologs of this clade, named MpPATH and MpPATI, appeared to be polarized when expressed in Arabidopsis roots (**Fig. 5J,K**). PAT18, also a clade VIII member, appeared to have lost the ancestral polarized localization and localized to the PM and endomembranes (**Fig. 5L**). We found that apart from clade VIII, PAT07 also localized to the PM (**Fig. 5M**), but in a non-polar way. PAT10 and PAT15 localized to the endomembranes with some PM signal (**Fig. 5N-O**), while PAT1, PAT17 and PAT23 localized only to endomembrane structures (**Fig. 5P-R**).

**Fig. 5.**
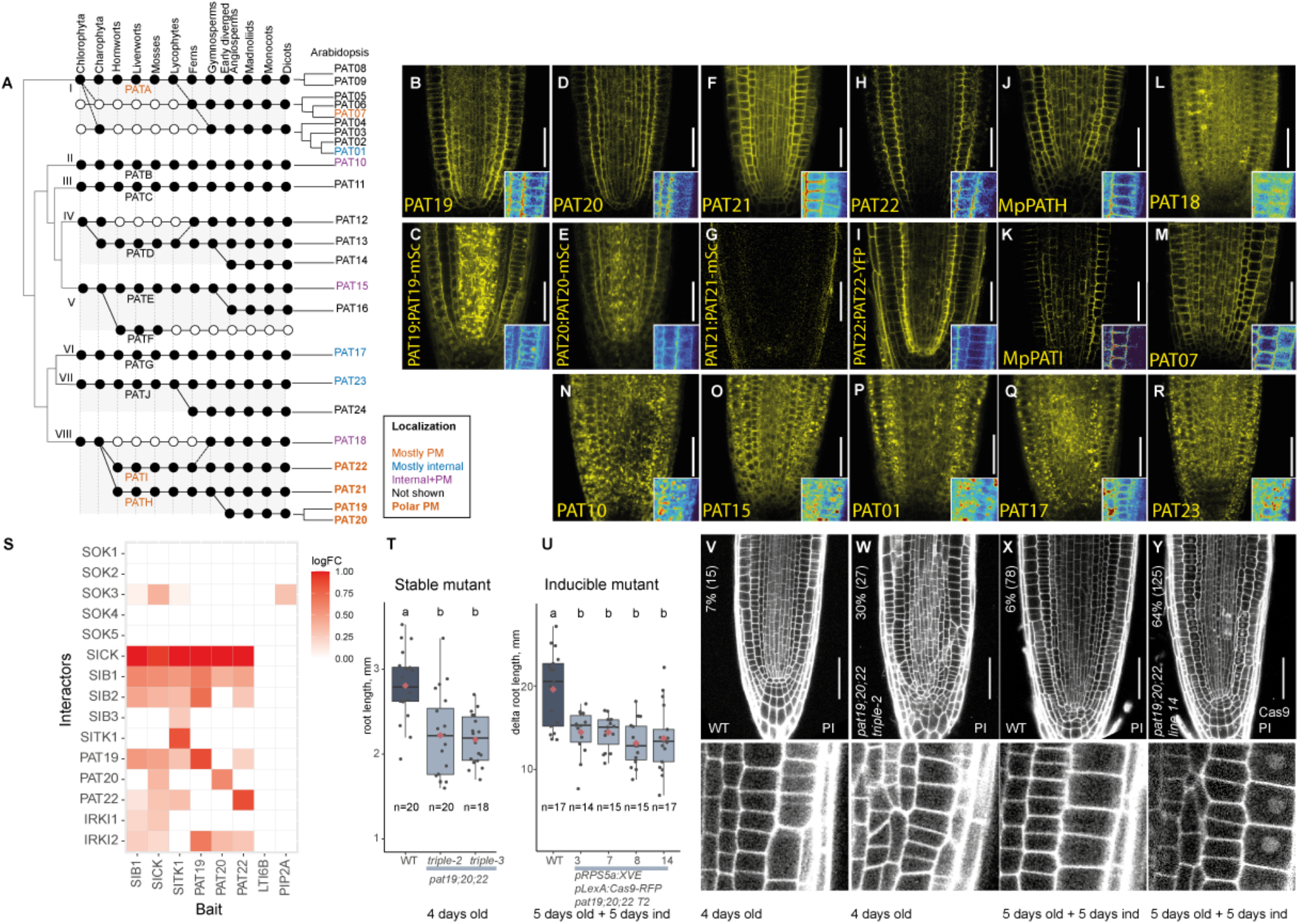
Identification of a polarized PAT subclade required for Arabidopsis root development. (**A**) Schematic phylogenetic tree showing presence/absence and orthology of PAT family members across key plant lineages. Black/white circles indicate homolog presence/absence; Marchantia homolog names are listed for each clade. Taxon order reflects the position of homolog splits and homolog number in the last common ancestor of the groups to the right, rather than evolutionary relationships. Black dashed lines denote hypothetical branches representing paraphyletic sequence groups. The right side shows sequence divergence within Brassicaceae, with Arabidopsis homologs. Roman numerals denote clades; color and font of Arabidopsis and Marchantia protein names indicate PAT localization. (**B**-**R**) Confocal images of roots of 7-days-old Arabidopsis seedlings expressing fusions of each indicated PAT (or MpPAT) to TbID-YFP (**B**,**D**,**F**,**H**,**J**,**K**,**L**,**M**,**N**,**O**,**P**,**Q**,**R**) from RPS5A promoter, or as mScarlet (**C**,**E**,**G**) or YFP (**I**) fusion to the native promoter. Scale bar: 50 µm. Bottom right corner of each image shows inset of a few epidermal and cortex cells with false color intensities. (**S**) Heatmap of normalized logFC values from Arabidopsis proximity labeling (lateral polar protein baits) for SOK lateral polar complex proteins. Newly generated PAT19, PAT20, and PAT22 bait data are shown alongside SIB, SICK, and SITK data from **Fig. 3**. All experiments used 7-day-old seedlings expressing pRPS5a:bait-TbID-YFP. (**T**) Boxplots of main root length in 4-day-old seedlings of two *pat19;20;22* alleles (and WT Col-0. Boxes show median (black line), 1st/3rd quartile (box borders), and 1.5x interquartile range (whiskers); red diamonds indicate mean, gray dots indicate individual data points. The number of observations per independent T2 line is shown below each box. Letters denote statistical groups (Tukey HSD post-hoc test). Raw data and p-values are in **Supplementary Data 4**. (**U**) Boxplots of main root length increase over induction days in four independent T2 inducible CRISPR *pat19;20;22* lines and non-fluorescent seed WT background. Boxplot elements, statistics, and data availability are as in (**T**). (**V**-**Y**) Confocal images of roots: (**V**) 4-day-old WT Col-0; (**W**) 4-day-old stable *pat19;20;22* mutant; (**X**) WT non-fluorescent seed plant grown 5 days on regular MS medium followed by 5 days on 17-β-estradiol plates; (**Y**) T2 inducible CRISPR *pat19;20;22* plant grown under the same conditions as (**X**). Roots stained with propidium iodide (PI); Cas9-RFP imaged in the same channel. Numbers indicate the percentage of roots with cell division defects and the total number of analyzed roots. Below each image, an inset shows normal (WT) or disturbed (mutant) cell division orientations.

We next asked whether all members of this group of polarized PAT enzymes are connected to the polar protein network. We generated TbID fusions to PAT19, PAT20 and PAT22 and performed proximity ligation experiments. We found all three PAT enzymes to associate with several members of the core polar network (SICK, SIB1, SIB2) as well as with IRKI (**Fig. 5S**). This suggests a shared role for PAT19, 20 and 22 in cell polarity in the root meristem.

Previously, mutants in these three PAT enzymes have been described, and a triple mutant displays developmental defects^48^. We confirmed that the stable *pat19;20;22* triple mutant has shorter roots (**Fig. 5T**). Furthermore, we generated inducible Cas9 lines to fully delete all three genes upon estradiol treatment (**Fig. S3A-B**), and confirmed that conditional deletion of PAT19, 20 and 22 shortened roots (**Fig. 5U, S4A-B**). Furthermore, we found that both stable and inducible *pat19;20;22* mutants had oblique cell division orientations in the root (**Fig. 5V-Y, Fig. S4C**), possibly indicative of defects in cell polarity. These phenotypes in the inducible CRISPR mutants were reproducible over 4-7 days of induction, even though they did not get more severe over time (**Fig. S4C**). The inducible deletion of an unrelated control gene PLP1 did not cause these phenotypes (**Fig. S3C, S4B-C**), confirming the specificity of the inducible gene editing. In summary, we identified a subfamily of polarized PAT enzymes that are required for normal growth and cell division in Arabidopsis roots.

### PAT enzymes are required for membrane targeting of SOK1

PAT enzymes target membrane localization of their substrates by transferring a palmitate to a cysteine. Several polar proteins, including SOK1 and SOK5, but also BRX and SGN1, contain cysteine residues essential for their polarity and PM association^29,36,49^. The discovery of polarly enriched PAT enzymes in the proximity of the SOK complex suggests that SOK itself or other complex members are substrates of these PAT enzymes.

SOK1 contains cysteine C233 essential for its PM association^29^. This cysteine is a part of an 80 amino acid polar domain identified in the SOK1-SOK2 domain swap experiment as being sufficient to target a SOK2 chimera to the SOK1 polar domain^28^. SOK proteins require a polymerization DIX domain for focused localization^29^. The human Dvl2 DIX domain is functionally equivalent to the *A. thaliana* DIX domain^29^ but is less likely to interact with other plant cell polarity regulators. We therefore used it as the polymerization module in place of the native domain, fusing the 80-amino-acid polarity domain to the Dvl2 DIX domain and mNeonGreen (**Fig. S5A**). This fusion, henceforth called miniSOK1, was sufficient to target the protein to the cell corner, where it localized in a manner similar to the full length protein (**Fig. S5B-C**). Thus, the polarity domain containing a critical cysteine contains all information for polar membrane targeting.

To address if SOK1 is acylated *in vivo*, we treated fluorescently labelled SOK lines with hydroxylamine (HA) that removes palmitoylation from cysteines and causes the PM-to-cytosol transition of membrane-anchored palmitoylated proteins^50^. HA treatment caused a substantial loss of PM association for full-length SOK1 in root cells (**Fig. 6A-B**). This effect was exaggerated in miniSOK1 roots (**Fig. 6C-D**), suggesting that S-acylation of the minimal targeting region is critical, but protein domains outside of this region also contribute to membrane interaction. In contrast, localization of the transmembrane PM markers NPSN12 and PIP1;4^51^ was unaffected by HA treatment (**Fig. 6E-F, Fig. S5D-E**), while LTI6B moved to intracellular structures (**Fig. S5F-G**), distinct from the cytosolic SOK signals.

**Fig. 6.**
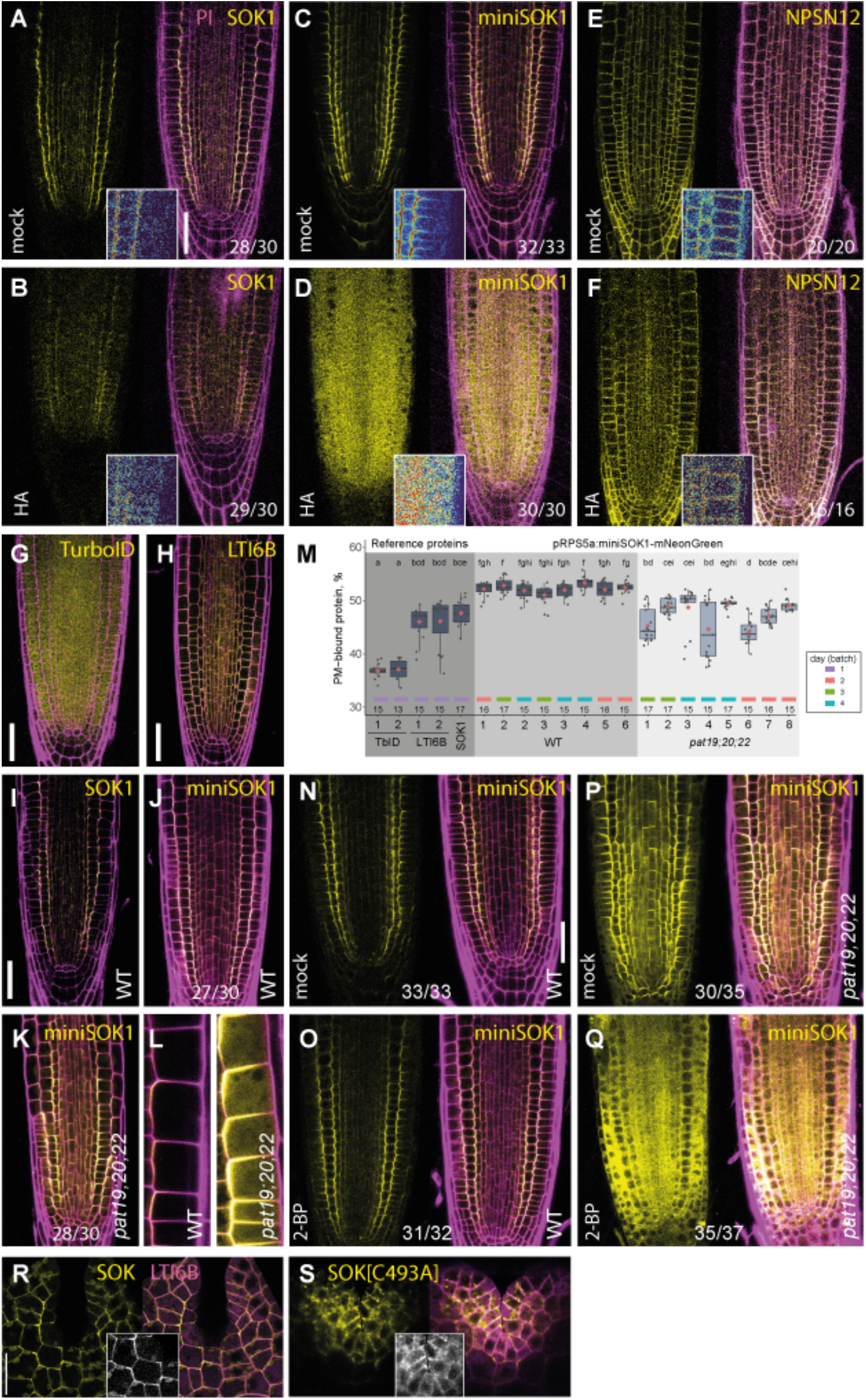
SOK1 PM association depends on the PAT19 subfamily of S-acylation enzymes. (**A**-**F**) Confocal images of 7-day-old roots expressing (**A**,**B**) pRPS5a:SOK1(full length)-YFP, (**C**,**D**) pRPS5a:miniSOK1-mNeonGreen, (**E**,**F**) pUB10:NPSN12-EYFP (WAVE131), treated for 4 h with 50 µM hydroxylamine (HA; **A**,**C**,**E**) or 50 µM NaCl (mock; **B**,**D**,**F**). Scale bar: 40 µm. Each panel shows the protein of interest (yellow) and a merge with propidium iodide (PI) cell wall stain (magenta); bottom right corners shows inset with false-color intensities. (**G**-**L**) Confocal images of 7-day-old roots expressing pRPS5a:TbID-YFP (**G**), pRPS5a:LTI6B-YFP (**H**), pRPS5a:SOK1(full length)-YFP (**I**) and pRPS5a:miniSOK1-mNeonGreen (**J**-**L**) in WT (**J**,**L**) or stable *pat19;20;22* mutant (**K**,**L**). (**M**) Quantification of the PM-associated fraction of fluorescent signal in all lines depicted in (**G-L**). Boxes show median (black line), 1st/3rd quartile (box borders), and 1.5x interquartile range (whiskers); red diamonds indicate mean, gray dots indicate individual data points. The number of observations per independent T2 line is shown below each box. Letters denote statistical groups (Tukey HSD post-hoc test). Imaging was done different experimental days in batches indicated by tiles below the boxes. Raw data and p-values are in **Supplementary Data 4**. (**N-Q**) pRPS5a:miniSOK1-mNeonGreen fluorescence in wild-type (**N**,**O**) and stable *pat19;20;22* mutant (**P**,**Q**) treated for 15 h with 20 µM 2-bromopalmitate (2-BP; **O**,**Q**) or equivalent DMSO (mock; **N**,**P**). Scale bar: 40 µm. (**R**,**S**) Confocal images of *Marchantia polymorpha* gemmae expressing pEF:LTI6B-mCherry PM marker (magenta) and pEF:MpSOK-TbID-YFP (**R**) or pEF:MpSOK[C493A]-TbID-YFP (**S**) (yellow). Insets show a few apical notch cells (SOK channel). Scale bar: 20 µm.

To directly test the involvement of PAT19/20/22 in SOK1 targeting, we introduced the miniSOK1 reporter, as well as reporters for SIB1 and SICK, into the stable *pat19;20;22* triple mutant (**Fig. 6G-M, Fig. S5H-M**). We first developed a method to quantify PM localization by measuring overlap with the Propidium Iodide cell wall stain, and benchmarked it with constitutively cytosolic (free TbID-YFP) and PM-localized (LTI6b) proteins (**Fig. 6M**). While polar membrane localization of all three proteins was retained in the mutant (**Fig. 6K,L; Fig. S5I,K**), the fraction of cytosolic signal was consistently increased for miniSOK1, but not for SIB1 or SICK (**Fig. 6M; Fig. S5L,M**), indicating a selective role for PAT19/20/22 in miniSOK1 targeting.

While this result indicated involvement of PAT19/20/22 in SOK1 membrane targeting, the effect was small, perhaps due to redundancy with other PAT enzymes. Given that several other PATs are PM-localized, and because the *pat19;20;22* triple mutant already suffers from fertility defects^48^, we chose a pharmacological approach to further test PAT involvement in SOK1 PM targeting. We used the stable triple mutant as a sensitized background to test the effect of the broad S-acylation inhibitor 2-bromopalmitate (2-BP). This compound inhibits all enzymatic pathways that utilize long chain acyl-CoA^52,53^. Because the triple mutant has reduced overall PAT activity, we reasoned it should be more sensitive to partial pharmacological inhibition than the WT, allowing us to detect an effect at a low, non-toxic 2-BP concentration.

In the WT background, 2-BP treatment did not cause any visible miniSOK1 localization changes (**Fig. 6N-O**), even though quantification showed a decrease in PM fraction, in line with the effect of HA treatment on miniSOK1 described above (**Fig. S5N**). In the triple mutant, the same concentration of 2-BP almost fully displaced miniSOK1 from the PM (**Fig. 6P-Q**). This result shows that the PAT19 subfamily is indispensable for stable PM association of SOK1, but other redundant enzymes contribute to the process.

While PAT19 homologs are present in the homologous lateral module in Marchantia, we also noticed that the cysteine-containing region is homologous between Arabidopsis and Marchantia SOK proteins, based on land plant SOSEKI phylogeny reported^29^ (**Fig. S5O**). Engineering a C493A point mutation in MpSOK led to significant loss of its polarity and PM association and higher protein accumulation in the cytosol (**Fig. 6R-S**), in line with Arabidopsis C233A mutant^29^. Overall, we showed the requirement for the S-acylation in a conserved protein motif of SOK proteins for their PM association.

### PAT19 is sufficient for SOK1 targeting to the plasma membrane

The finding that SOK1 PM localization depends on palmitoylation in a PAT19/20/22-dependent manner suggests a simple model for polar membrane targeting in which polarized PAT enzymes facilitate localized insertion of SOK1 into the PM. This minimal targeting model would rely on this enzyme-substrate interaction being sufficient for membrane insertion. We used two heterologous model systems to test the sufficiency of the PAT19-SOK1 interaction.

First, we expressed Arabidopsis SOK1 as a fluorescent fusion in human HeLa and U2OS cells and found the protein to be present in the cytoplasm (**Fig. 7A-D, S6A-N**). In maximum intensity projections, the brightest signal was observed in the perinuclear region, where the cell is thickest, consistent with the protein accumulating in the cytosol and being excluded from the nucleus (**Fig. 7A, S6F**). Arabidopsis PAT19 localized in these cells to PM and Golgi (**Fig. 7B, S6B,D,G,K,M**), in line with native plant localization. When PAT19 was co-expressed, we observed a different pattern of SOK1 localization, where the bulk cytoplasmic signal was lost and the protein was evenly distributed over the cell surface, consistent with PM targeting (**Fig. 7B, S6G**). We engineered either a mutation in the potential palmitoylation substrate cysteine in SOK1 (C233A or C233S), or in the catalytic site of PAT19 (C204S) (**Fig. S6A,C,E,J,L,N**). All of these mutations led to the loss of the effect of PAT19 on SOK1 localization (**Fig. 7C-D, S6B,H,I,K**), consistent with sufficiency of PAT19-dependent SOK1 palmitoylation for PM targeting. However, since adherent cells are thin, it was difficult to quantify protein partitioning between the cytosol and the PM. We therefore next turned to the fruit fly (Drosophila melanogaster) wing epithelium as a model. This tissue has been widely used to study polar membrane targeting^54,55^, and the morphology of epithelial cells allows clear visualization of both the PM and cytosol in z-sections (**Fig. 7E**). We ubiquitously expressed a SOK1-mEGFP fusion protein in prepupal or early pupal wings and found that it showed some membrane association, and a large cytosolic fraction represented by punctate structures (**Fig. 7F, S6O**). Co-expression of 3xHA-tagged PAT19 caused a strong reduction of SOK1 cytosolic puncta signal, that could be explained by translocation of SOK1-mEGFP to the plasma membrane or by PAT19 activity either promoting puncta degradation or reducing SOK1 aggregation (**Fig. 7G, K-M, S6P**).

**Fig. 7.**
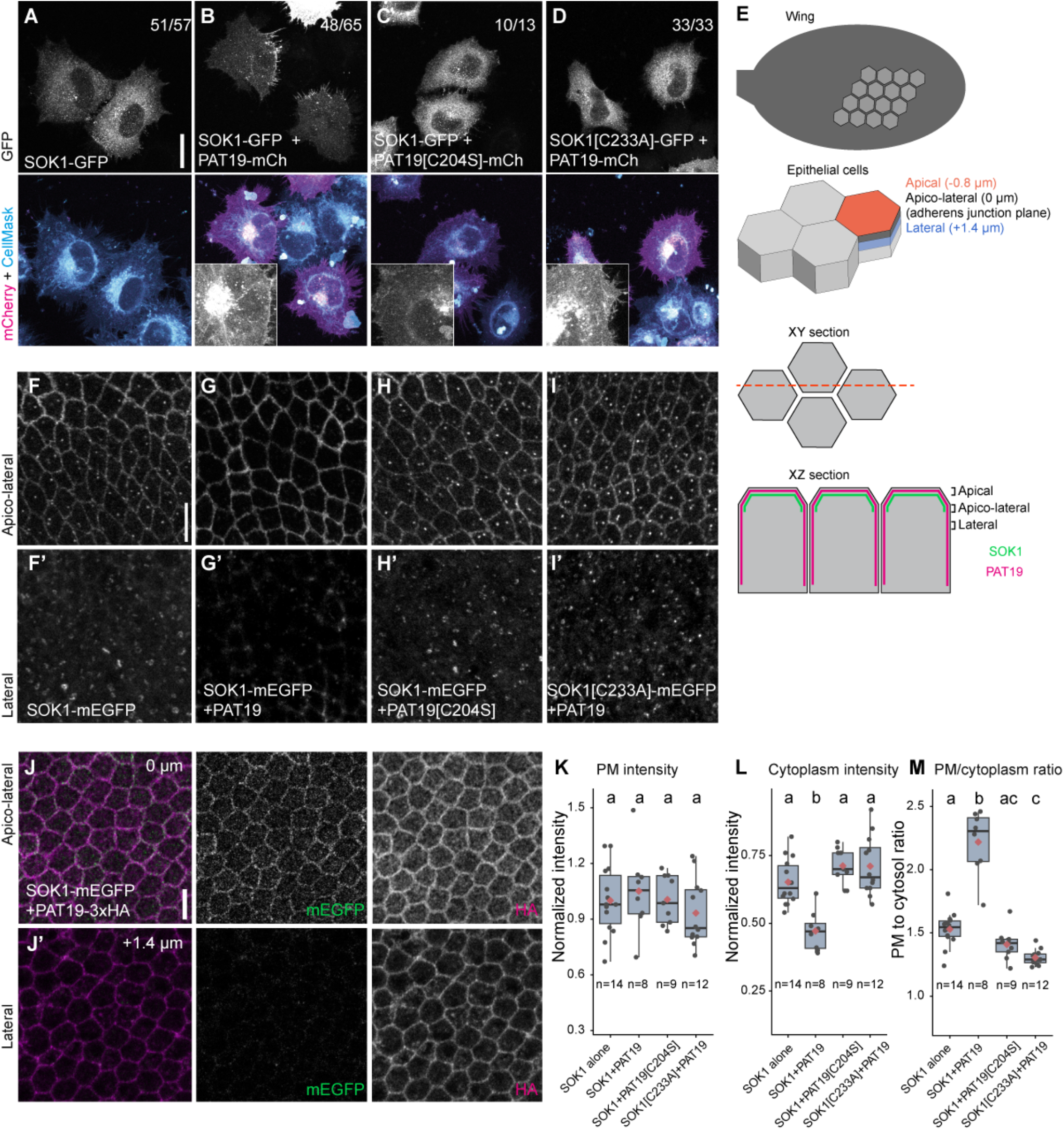
PAT19 promotes plasma membrane association of SOK1 in HeLa cells and Drosophila epithelia. (**A**-**D**) Maximal intensity projections of confocal images of fixed HeLa cells transiently expressing combinations of full-length SOK1-GFP, PAT19-mCherry, and their cysteine mutants (SOK1[C233S/A], PAT19[C204S]). Membranes were stained with CellMask Deep Red. Insets show a fragment of a cell with PAT19-mCherry. Numbers indicate the number of cells showing the SOK1/PAT19 localization pattern in the representative image, across 2 independent experiments. Scale bar 20 µm. (**E**) A scheme of confocal section planes in Drosophila wings. At the top is an XY section, in which the images look down onto the apical wing surface. The red line indicates the position of an example XZ section below, showing the apical domain, the apico-lateral junctional domain and a lateral region. (**F**-**I**) mEGFP fluorescence in 6 hr after puparium formation (APF) live prepupal wings expressing SOK1-mEGFP (**F**), SOK1-mEGFP and PAT19-3xHA (**G**), SOK1-mEGFP and PAT19[C204S]-3xHA (**H**) or SOK1[C233A]-mEGFP and PAT19-3xHA (**I**). Top panels are an average projection of three 0.2 µm confocal slices, centered around the region of maximal SOK1-mEGFP plasma membrane signal, at the apicolateral junctional region. Lower panels are an average projection of ten lateral 0.2 µm confocal slices, starting 0.6 µm more basally. Scale bar 5 µm. (**J**) mEGFP fluorescence (green) and HA immunolabelling (magenta) in a 28 hr APF fixed pupal wing expressing SOK1-mEGFP and PAT19-3xHA. Confocal section at the level of maximal apico-lateral SOK1-mEGFP plasma membrane signal. (**J’**) is a confocal section that is 1.4 µm more basal. Scale bar 5 µm. (**K-M**) Quantification of membrane intensity (**K**), cytoplasm intensity (**L**) and membrane/cytoplasm ratio (**M**) of SOK1-mEGFP fluorescence for the genotypes in (**E-I**), normalized to mean SOK1-mEGFP membrane intensity in the absence of PAT19. Plasma membrane (PM) intensity, cytoplasm intensity and PM-cytoplasm ratio are measured from an average projection of the apico-lateral region in (**E-I**) plus the next seven more basal slices. Boxes show median (black line), 1st/3rd quartile (box borders), and 1.5x interquartile range (whiskers); red diamonds indicate mean, gray dots indicate individual data points. The number below each boxplot indicates sample size (number of flies). Statistics are ANOVA with Tukey’s multiple comparisons test, letters denote statistical groups. Raw data and p-values are in **Supplementary Data 4**.

Interestingly, SOK1-mEGFP was largely excluded from lateral plasma membrane domains, but instead localized specifically to apical and apico-lateral regions of the plasma membrane (**Fig. 7E,J, S6P**). In contrast, immuno-labelling of PAT19-3xHA showed that it was distributed throughout the plasma membrane (**Fig. 7J**). An increase in plasma membrane localization of SOK1-mEGFP would be expected if PAT19 promoted translocation of SOK1-mEGFP to the plasma membrane. This was not evident in the quantitation; however the population of SOK1-mEGFP that localized to the apical plasma membrane surface was not captured in these data. Consistent with PAT19 enzymatic activity acting on C233 in SOK1, catalytically dead PAT19 did not change SOK1 localization and C233 mutant SOK1 localization was also unaltered by PAT19 co-expression (**Fig. 7H-I, K-M**).

We conclude that PAT19 catalytic activity is sufficient to target SOK1 to the plasma membrane by palmitoylation in two heterologous systems in the absence of any other plant proteins.

## Discussion

Cell polarity appears to be a major driving force in the evolution of complex tissues in land plants, yet many plant polarity proteins and their functions are not conserved among land plants^5,12,56^. This may reflect a gap in our understanding of land plant cell polarity rather than the existence of many independent polarity pathways, and suggests that most currently known polarity proteins are “clients” rather than regulators of polar targeting. SOK proteins are conserved throughout land plants^28,29^, and their edge localization may provide a missing link between cell geometry and intracellular processes^57^. Using cross-species interactomics, we identify key components of this polarity pathway and show that SOK proteins are part of a conserved and polarly localized protein network.

This conserved lateral polar protein complex contains PAT enzymes that are biased toward the inner PM face in Arabidopsis roots. These PATs form a monophyletic clade that arose in the land plant common ancestor. Whether the polarity of this clade is an ancestral trait remains unclear, as its Marchantia orthologs are evenly distributed at the PM in gemmae. Conversely, MEE59 shows the opposite pattern - polar in Marchantia but not in Arabidopsis. This may reflect differences between orthologs, or between tissues, since Marchantia gemmae are photosynthetic gametophyte tissue whereas Arabidopsis tissues studied here are sporophyte root tissue. Still, the tendency of Marchantia PATH and PATI to localize polarly in Arabidopsis roots suggests a conserved, root-specific polarization mechanism - potentially linked to distinct properties of this tissue - that biases PAT19 clade enzymes toward the endodermis-cortex junction. This is supported by a similar localization bias among other Arabidopsis polar proteins^28,49,58^.

PATs mediate membrane targeting of proteins, and we propose that their polar localization maintains an asymmetric protein composition across the cortical cell region. We identify SOK1 as one candidate polar substrate of Arabidopsis PAT19. S-acylation is required for SOK1’s PM association across the land plant lineage, but is likely not the only PM targeting mechanism, as full-length Arabidopsis SOK1 is less sensitive to HA treatment than miniSOK1. In Drosophila cells, SOK1 partially localizes to the PM even without plant PAT, suggesting alternative PM-targeting mechanisms such as electrostatic interactions with phospholipids. Notably, SOK1 is polar in Drosophila cells while PAT19 is not, pointing to additional, PAT-independent polarization mechanisms. Experiments with *pat19;20;22* Arabidopsis mutants suggest SOK1 is predominantly, but not exclusively, targeted to the PM by the PAT19 subclade. Even in better-studied animal systems, how enzyme-substrate specificity is achieved among PATs remains unclear: limited in vitro assays on human PATs indicate that some enzymes have preferred substrates, with residues flanking the target cysteine affecting reaction efficiency^59^, while other substrates can be palmitoylated by multiple PATs, even across species^60,61^. In Arabidopsis, several PM-localized PAT subclades may compensate for PAT19 loss in the *pat19;20;22* mutant. Further in vitro assays on plant PATs will be needed to define the biochemical basis of substrate recognition. PAT19 substrates are also not limited to polar proteins, as shown by the nonpolar substrate BSK1^48^; whole-proteome analysis of S-acylation and subcellular localization in the mutant could help distinguish polarity-specific from more general functions of the PAT19 subclade.

We observed occasional cell division orientation defects in the stable *pat19;20;22* mutant, which were exacerbated in the inducible line - likely because genetic redundancy allows for compensation from the early developmental stages in the stable mutant, whereas sudden loss of PAT19 in the inducible line does not. Incomplete penetrance may reflect stochastic activation of compensatory mechanisms and/or cell-type-specific differences. These phenotypes suggest that the lateral polar complex contributes to oriented cell division control in the root apical meristem, though not necessarily through SOK1 directly. Since the functions of SOK and other lateral module components (SIB, SICK, SITK) remain unstudied, these or other PAT substrates may underlie the observed division orientation phenotype. SITK, homologous to the IRK/PXC2/KOIN receptor-like kinases that control root meristem cell divisions^58,62^, is a promising candidate linking these functions.

Finally, our cross-species interactome comparison suggests a small conserved core of polar protein interactors surrounded by a larger, more divergent set reflecting diverged protein functions across species. More broadly, the foundations of cell polarity may have been present in the last eukaryotic common ancestor, with different eukaryotic lineages later building distinct polarity systems and multicellular bodies from the same basic components. DIX-domain proteins, for instance, contribute to polarity in three independently multicellular eukaryotic lineages — Metazoa^63^, TSAR^64^, and land plants^29^, even though the rest of the protein sequences, including S-acylation sites, is not conserved across kingdoms; DIX domain polymerization itself may underlie polarity by restricting protein diffusion. Similarly, polarized PAT-mediated PM targeting may serve as a general mechanism for concentrating proteins at the membrane and generating cellular asymmetry. Acylation is a common post-translational modification among polar proteins beyond plants: Rho GTPases (ROP in plants, Cdc42 in animals and yeast) use S-acylation for PM localization during polar growth^65,66^, and in mammalian epithelial cells, polarly localized DHHC PATs regulate protein composition at tight junctions67. Restricting enzyme activity to concentrate a substrate may thus be a general cell polarity mechanism: even where individual proteins are not homologous across kingdoms, they may share conserved underlying principles of polarity.

## Acknowledgements

We are grateful to Yan Zhang and Fei Liu for sharing stable *pat19;20;22* mutants seeds, Hirokazu Tsukaya for 35S:AN-GFP and Marie-Cécile Caillaud for *sac9* pSAC9:mCitrin-SAC9 lines, Sjef Boeren for support with mass spectrometry, Arjen Bader for support with imaging and Joris Sprakel for advice on image analysis.

This paper was typeset with the bioRxiv word template by @Chrelli: www.github.com/chrelli/bioRxiv-word-template

## Author contributions

Conceptualization: EMP, VJ, CA, DW

Data Curation: EMP, MR, SM

Investigation: EMP, CA, MvD, VJ, MR, JJR, SM, CS, HS, JCMM

Formal analysis: EMP, CA, MvD, VJ, MR, JJR, SM, CS, HS, JCMM, AA, DS, DW

Funding Acquisition: MvD, AA, DS, DW

Supervision: CA, MR, AA, DS, DW

Validation: EMP, CA, MvD, MR, CS, HS, JCCM

Visualization: EMP, CA, MvD, MR, SM, CS, HS, JCCM

Project Administration: EMP, CA, DW

Writing – Original Draft: EMP, DW

Writing – Review & Editing: All authors

## Data availability

Raw proteomics data is available in MassIVE (MSV000102532) and PRIDE (PXD081338).

Shiny app for proximity labeling data: https://weijerslab.shinyapps.io/polar_protein_proxitome/

Shiny app code : https://github.com/evgeniyampukhovaya/polar_protein_proxitome

PAT phylogeny: https://zenodo.org/records/21411520

Code for protein PM fraction quantification in Arabidopsis roots: https://github.com/evgeniyampukhovaya/PM_signal_quantification_in_Arabidopsis_roots

## Materials and Methods

### Plant material and growth conditions

*Arabidopsis thaliana* wild type ecotype Columbia-0 (Col-0) and all Arabidopsis mutants and transgenics were grown on vertical square Petri plates on ½ strength Murashige and Skoog (MS) basal medium (Duchefa M0221), pH 5.8, supplemented with 0.8 % plant agar (Duchefa P1001) and 1 % MES. Seeds were sterilized with 75% ethanol/25% bleach for 3 min, washed 3 times with 70% ethanol and once in 96% ethanol, dried in the flow cabinet for 2-3 h, mixed with sterile milliQ water or 0.1% plant agar and distributed on the plate agar surface, then vernalized at 4°C for 1-4 days. Plants were cultured under 90-100 µmol photons m-2 s-1 white light in long day conditions (16 h light/8 h dark cycle) at 22°C and 75% humidity. pSOK1:SOK1-YFP, pSOK2:SOK2-YFP, pSOK3:SOK3-YFP, pRPS5a:SOK1-YFP, pRPS5a:SOK2-FP, pRPS5a:SOK3-YFP^1^, pAN:AN-TdT^2^, p35S:AN-GFP^3^, *sac9* pSAC9:mCit-SAC9^4^, pUB10:NPSN12-EYFP (WAVE131), pUB10:PIP1;4-EYFP (WAVE138)^5^, *pat19;20;22* stable mutant^6^ were described before. Generation of the rest of the transgenic lines is described below and in Supplementary data 5.

*Marchantia polymorpha* Takaragaike-1 (Tak-1) was used as the wild-type accession for all Marchantia lines and experiments. Plants were cultured on ½ strength Gamborg B5 medium (Duchefa G0209) at 22°C under constant white light (40 µmol photons m^−2^ s^−1^).

### Plasmid construction for plant overexpression and reporter lines

Primers, backbone vectors and resulting Arabidopsis transgenic lines are listed in Supplementary data 5. Vector backbones for proximity labeling, over-expression and reporter lines were based on pPLV backbones described before^7^. For the initial set of TurboID lines (Fig. 2), the bait CDS were amplified from Arabidopsis cDNA and cloned into a vector VJp016 (pRPS5a:LIC:TurboID-sYFP2, phosphinothricin (PPT) selection) digested with HpaI (NEB R0105S) using SLiCE method^8^. For the candidates from the dataset (Fig. 3) and for PAT lines (Fig. 5), the protein CDS was amplified from Arabidopsis gDNA using Phusion-Flash High Fidelity PCR Mastermix (Thermo F548L), purified from agarose gel using NucleoSpin Gel and PCR Clean-up kit (Macherey-Nagel 740609.50) and cloned into a vector EPp022 (pRPS5a:LIC:TurboIDs-YFP2, FastRed selection) digested with Eco105I FastDigest (Thermo FD0404) using NEBuilder® HiFi DNA Assembly Master Mix (NEB E2621S). Marchantia PATI and PATH for Arabidopsis expression were amplified from Marchantia cDNA and cloned into EPp022. For native promoter reporters, genes and ∼2.5 kb upstream promoter regions were amplified from Arabidopsis gDNA and cloned into pPLV17 (C-terminal YFP fusions) or similar custom-made pPLV backbones for mNeon-Green, mTurquoise2 and mScarlet fusions as described for TurboID vectors. For miniSOK1 reporter cloning, hDvl2-DIX domain from the human Dvl2 protein was amplified from a construct described in^2^, fused to 80 amino acid part of SOK1 CDS and cloned into custom made pRPS5a:LIC-mNeonGreen pPLV-based vector.

For Marchantia plasmid generation, genomic fragments of *MpSOK, MpSIB, MpSICK* and the rest of the candidates (Fig. 4) were PCR-amplified from *Marchantia* genomic DNA (primers listed in Supplementary data 5). Amplicons were inserted into linearized pDONR221 (linearized using primers CA1500/CA1501) by NEBuilder® HiFi DNA Assembly (New England Biolabs) to generate Gateway entry clones. Entry clones were recombined into a modified pMpGWB destination vector carrying an N-terminal (pMpGWB101) or C-terminal (pMpGWB103) TurboID-YFP cassette via LR reaction using Gateway® LR Clonase® II Enzyme Mix (Thermo Fisher Scientific 11791020), yielding fusion constructs.

*E. coli* strain DH5α electrocompetent cells were used for plasmid isolation.

### Plant transformation

All the constructs were sequenced and transformed into wild type (WT) Arabidopsis (Col-0) by floral dip^9^ using *Agrobacterium tumefaciens* strain GV3101:pMp90 (pSoup). Transformants were selected on ½ MS medium with 0.8% Plant agar, 1% MES and 15 mg/L phosphinothricin (PPT) or 50 mg/L kanamycin, or by selecting red/green fluorescent seeds on Leica M205 FA stereomicroscope. Stable mutant *pat19;20;22*^6^ was transformed with miniSOK1, SICK and SIB1 reporters in a similar way.

Marchantia Tak-1 spores were transformed with the above constructs using *Agrobacterium tumefaciens* strain GV3101:pMp90, following the protocol^10^. Transformants were selected on ½ strength B5 medium supplemented with 10 mg/L hygromycin and 100 µg/mL cefotaxime.

### IP-MS experiments in Arabidopsis

IP-MS experiments for SOK1-3, AN, DLC1 and SIB1 (Fig. 1A-F) were done according to the protocol described in^2^. For the SICK IP-MS experiment (Fig. 1G), seeds of three independent segregating T3 lines of pRPS5a:SICK-TbID-YFP in were mixed in equal proportions; Col-0 WT seeds were used as a control. Seedlings were grown for 7 days under long day light conditions; then roots from 30 plates for each genotype were cut with scalpel, ground in a mortar, and frozen in liquid nitrogen. Each genotype had 3 technical replicates from the same lysate generated from the same pool of tissue powder.

Root powder was diluted 1:1 volume ratio with the cold lysis buffer (50mM Tris pH 8.0, 150mM NaCl, 2mM MgCl2, 1mM EDTA, 1mM DTT, 0.2% NP40, 10% Glycerol, 1x cOmplete Protease Inhibitor Cocktail (Roche 04693116001)), mixed and sonicated in the precooled water bath 3 times at 15 s at 80% amplitude. After that, it was centrifuged 30 min at 21 000 g 4°C. Protein concentration in the lysates was determined with Bradford protein assay (Bio-Rad 5000006), 1 mg of total protein in the total volume of 1 ml was incubated with 50 µl ChromoTek GFP-Trap® Agarose (proteintech gta) equilibrated with the lysis buffer. Samples were incubated for 90 min at 4°C rotating, then unbound proteins were washed 3 times with the lysis buffer, 3 times with the same buffer without detergent, and 3 times with 50 mM ammonium bicarbonate. Free cysteine alkylation was performed for 30 min at RT in 20 mM iodacetamide in 50 mM ammonium bicarbonate in 1 ml total volume; then the beads were mixed with 0.25 µg of trypsin (Roche 11047841001) per sample in the total volume of 100 ul 50 mM ammonium bicarbonate. Trypsin digestion was performed overnight at room temperature, then the peptides were acidified with 10% trifluoroacetic acid to pH ∼ 2-3 checked with the indicator paper. C18 columns made in 200 µl pipette tips with LiChroprep RP-18 (25-40 µm) (Merk 1.09303) and Empore SPE disks (Sigma 66889-U) were pre-washed once with 50 µl 80% acetonitrile 0.1% formic acid and once with 50 µl 0.1 % Formic acid. Peptides were loaded on C18 columns, washed 2 times with 50 µl 0.1 % Formic acid, eluted with 30 µl 80% acetonitrile 0.1% formic acid, vacuum-dried and diluted in 25 µl 0.1 % Formic acid.

Peptides were separated on reversed-phase nano LC (Thermo Vanquish Neo) with 30 minute gradient and MS/MS spectra were measured on an Orbitrap Exploris 480 (Thermo) in DDA mode, as described^11^. Spectral search and label-free quantification (LFQ) were performed using MaxQuant software^12^ using UniProt Araport11 Arabidopsis reference proteome (UniProt ID UP000006548), with the parameters described in^11^. To get the statistical significance enrichment values from MaxQuant LFQ values protein groups outputs of all the IP-MS experiments (Fig. 1A-G), Perseus software^13^ was used. Data was filtered for reverse, contaminants, and only identified by site protein groups. Intensity values were log2 transformed and filtered to contain at least 2 valid values in each group. Remaining missing values were imputed from a normal distribution using standard settings in Perseus (width: 0.3, down shift: 1.8). Three technical replicates from pRPS5a:SICK-TbID-YFP samples were compared with WT Col-0 samples using 2-sample Student T-test. Benjamini-Hochberg FDR on Perseus-derived p values was calculated using R v.4.4.2 (R Core Team, 2024) (FDR = p.adjust(p_val, method = ‘BH’)). R package UniProt.ws was used to match Uniprot and gene IDs. Volcano plot visualization was made in R ggplot2 package.

### TurboID proximity labeling – sample preparation

Proximity labeling experiments on Arabidopsis (Fig. 2, 3, 5S) were performed in T3 or T4 generation segregating (non-homozygous) lines without selection for the presence of the transgene. Seeds were sterilized as described above, diluted with 0.1% plant agar and incubated for 1 day in the dark at 4°C. Vernalized seeds were sown on ½ MS 0.8% plant agar, 1% MES plates on a sterile nylon mesh in 3 rows and grew vertically for 7 days under long day light conditions. On the 7th day, the plates were flooded with liquid ½ MS medium containing 50 µM biotin (Duchefa B0603), 1% MES (pH 5.7), and incubated horizontally for 6 or 24 h. After that, the roots or whole seedlings were harvested in liquid nitrogen, grinded with mortar and pestle and stored at -80^°^C for further processing.

For Marchantia TurboID experiments (Fig. 4), gemmae were densely spread on B5 medium plates covered with a nylon mesh and grown for 10 days prior to biotin treatment. After 10 days of growth, young thalli were treated with 50 µM water biotin solution for 24 h to induce proximity biotinylation.

The sample preparation protocol was modified from^14^. Ground plant tissue powder from 35 square Petri plates (Arabidopsis roots), 20 plates (Arabidopsis whole seedlings) or 5 plates (Marchantia thalli) (or equivalent to ∼25 ml tissue powder in 50 ml falcon tube) was mixed with 1:1 volume of RIPA buffer (50 mM Tris pH 8.0, 150 mM NaCl, 2 mM MgCl2, 1 mM EDTA, 1 mM DTT, 1% NP40, 0.5% Sodium Deoxycholate, 10% Glycerol, 1x cOmplete Protease Inhibitor cocktail (Roche 04693116001)), sonicated in 4°C water bath (Qsonica) at 90% amplitude and centrifuged at 50000 g for 30 min. The protein extract was filtered from excess biotin through 10 kDa Amicon filters (Sigma UFC5010). Protein concentration was measured with Bradford protein assay (Bio-Rad 5000006). The same lysate was used for 3 technical replicates. An equal total protein amount in each experiment (1-5 mg of total protein, see Supplementary data 2&3 for details) in the total volume of 500 µl was mixed with 50 µl of Streptavidin Magnetic Sepharose beads (Cytiva 28985799) and incubated overnight at 4°C rotating head-over-tail. Second day, the beads were washed from unbound protein 3 times with 1 ml RIPA buffer, 3 times with 1 ml RIPA buffer without detergent and 3 times with 1 ml 50 mM ammonium bicarbonate, alkylated with 50 mM acrylamide in 50mM ammonium bicarbonate for 30 min at room temperature and incubated with 0.5 ug of trypsin (Roche 11047841001) per sample in 100 µl 50 mM ammonium bicarbonate overnight. C18 peptide purification was done on the third day as described for IP-MS samples above.

### TurboID proximity labeling – data analysis

MS measurement, raw spectra analysis and statistical comparison of LFQ values was performed as described for IP-MS experiments above. Transgenic lines expressing bait polar proteins were compared against control line expressing the empty TurboID-YFP vector (free ligase control). Big screen and biological replications were done in batches (Supplementary data 2&3), each batch including its own control.

Datasets were filtered for Benjamini-Hochberg FDR < 0.05 and log2FC>0 or log2FC>1 using R v.4.4.2 (R CoreTeam, 2024). Then logFC values were divided by the highest significant logFC in each dataset to normalize them in a range between 0 and 1, and all the datasets were merged in a wide format, with zero values indicating absence of interaction. It was used to filter for the proteins identified in a higher number of experiments, and to visualize interaction profiles of the proteins of interests – heatmaps depicting normalized logFC in all the datasets, using R ggplot2.

K-means clustering was performed in R v.4.4.2 using kmeans() function on a wide format file with normalized log2FC values. Optimal cluster number was defined by the elbow method as following:

~~~
iner%a <-vector()
for (i in 1:30) {
 kmeans_model <-kmeans(wide_df_for_clustering, centers = i)
 iner%a[i] <-kmeans_model$tot.withinss
}
p <-ggplot(data.frame(Clusters = 1:30, Iner%a = iner%a), aes(x = Clusters, y = Iner%a)) +
 geom_line() +
 geom_point() +
 labs(%tle = “Elbow Method for Op%mal Cluster Number”) +
 theme_bw()
~~~

For conservation analysis between Marchantia and Arabidopsis datasets (Fig. 4D), all the interactors were assigned to orthogroups as described in^11^. Orthogroups were matched to gene ontology (GO) terms based on the Arabidopsis orthologs using UniProt.ws R package, and the orthogroups containing GO terms matching regular expressions “nucle|ribosom|chloroplast|mitochondr” (cellular compartment), “translat|chlorophyll|metabolic process|glycolytic” (biological process) were excluded. The remaining orthogroups for Marchantia and Arabidopsis were considered as specific interactors and intersected to make a Venn diagram.

All the original unfiltered proximity labeling datasets were imported in R Shiny app. R packages tidyverse, shiny and reactable were used for data visualization and filtering.

### Phylogenetic analysis of PAT proteins

PAT phylogeny (Fig. 5A) was built according to a previously published pipeline^15^. Reference PAT sequences from *A. thaliana, A. thricopoda, P. patens, M. polymorpha, S. lycopersicum, M. truncatula, Z. mays, M. biondii, S. moellendorffii* were collected from Plaza database^16^. They were aligned with Clustal Omega^17^, using which a HMM was built that was used for a search in HMMER against One thousand plant transcriptomes (OneKP) dataset^18^. Significant hits were used for reciprocal Blast against Araport11 *A*.*thaliana* genome to confirm the orthologous relationship. Sequences for the phylogeny construction were aligned using ‘genafpair’ algorithm in MAFFT^19^, positions with large gaps (>90%) were removed with TriMal^20^. The phylogenetic tree was built using Maximum Likelihood (ML) method in IQ-Tree^21^ with 1000 rapid bootstraps. The resulting tree of 2972 sequences from 105 species was analyzed in iTOL^22^ using species classification according to the major Viridiplantae lineages. The presence and absence of PAT orthologs across different lineages was plotted in Fig. 5A using the iTOL tree as a reference.

### Arabidopsis CRISPR lines – DNA constructs

For inducible gene knock-out, an inducible CRISPR system^23^ was used. Single guide RNA (sgRNA) sequences for *PAT19, PAT20, PAT22, PLP1* exons were designed using the CHOPCHOP web tool^24^. Complementary forward and reverse primers (Supplementary data 4) carrying sgRNA sequences and ligation adapters were annealed by heating to 95°C and cooling to RT and introduced to pRU41-44 vectors^25^ digested with BbsI-HF (NEB R3539S) using T4 DNA ligase (NEB M0202T). Correct clones were verified by Sanger sequencing using oRU385 primer. The resulting plasmids were introduced into pSF464 first intermediate vector^25^ by GoldenGate reaction using T4 DNA ligase (NEB M0202T) and BsaI-HFv2 (NEB R3733S). *E. coli* strain DH5α was used for cloning and strain DB3.1 was used for backbone vector propagation. The assembly was confirmed by colony PCR (OneTaq 2X MasterMix, NEB M0482S). The 4 sgRNA-cassette was then amplified with primers oCS017, oCS018, the PCR fragment was gel-purified. The second intermediate vector p2R3z-BASI-ccdBBSAI (Addgene #118389^23^) was digested with BsaI-HFv2 (NEB), the 2.2 kb product was gel-purified. The 4 guides PCR fragment was combined with p2R3z-BASI-ccdB-BSAI fragment in a cloning reaction with NEBuilder® HiFi DNA Assembly Master Mix. The resulting pAtU3/6-sgRNA entry vector was confirmed by Nanopore sequencing and used as a box3 vector in a multiple Gateway LR reaction using Gateway® LR Clonase® II Enzyme Mix (Thermo Fisher Scientific 11791020). The other reaction components were pRPS5a:XVE_pLexA vector (box1^26^), Cas9p-tagRFP (box2, Addgene #118386), pFG7m34GW (Addgene #133747) destination vector. The final product was transformed into *A*.*thaliana* ecotype Col-0 WT.

### Arabidopsis CRISPR lines – Line selection and imaging

Fluorescent T1 transformant seeds and non-fluorescent seeds from the same batch (WT control) were grown for 5 days under long day conditions. Seedlings from fluorescent and control seeds were transferred in the sterile hood to ½ MS plates supplemented with 5 µM 17-β-estradiol (Sigma E2758). After 5 days of induction, seedlings were placed in the droplet of 20 µg/ml Propidium Iodide (PI) water solution and imaged with a Leica SP8 multiphoton (MP) microscope (see Confocal miscroscopy section below). After imaging, T1 seedlings were transferred to soil and further used for genotyping (described below) and T2 generation analysis.

For T2 (WT background), 4-5 T1 lines per construct showing both editing and phenotype/Cas9-RFP signal in T1 experiments were chosen. Seeds were selected, sterilized and grown as described for T1, non-fluorescent WT control seeds were selected from the same segregating T2 lines and treated same way as fluorescent seeds. After 5 days of growth, 15-20 seedlings per line were transferred to the ½ MS plates supplemented with 5 µM 17-β-estradiol. Plates were scanned with Epson Perfection V550 Photo scanner (day 0) and then continued growth under long day conditions. After 4, 5, 6 or 7 days of induction (new plate for each time point), plates were again scanned, roots were imaged with PI stain under Leica SP8 MP microscope. Main root length was measured from plate scans in ImageJ with Simple Neurite tracer plugin_27_. The increase in root length (delta root length) was quantified as a difference between the length of the same root before and after induction; these values were compared between independent T2 lines and non-fluorescent control using aov (delta_root_length ∼ Genotype) and TukeyHSD functions in R v4.4.2. Increases in growth values were visualized using R ggplot2, groups of significant difference were calculated with multcompView R package. Microscopy images of root apical meristem were processed using ImageJ and manually classified into having or not cell division defects and Cas9-RFP signal.

### Arabidopsis CRISPR lines – Genotyping

For T1 genotyping, the seedlings after confocal imaging were transferred to soil. After 2-3 weeks, young leaves were collected and placed on top of ½ MS 0.8% plant agar plates supplemented with 5 µM 17-β-estradiol for 2 days under long day light conditions. The leaf pieces were collected using a Unicore FTA Sample Punch Kit (Qiagen WB100028), DNA samples from the FTA card were used for PCR genotyping using OneTaq 2x mastermix (NEB M0482S). Genotyping primers are listed in Supplementary data 5. In case of single-guide mutations (*PAT20, PAT22*), PCR fragments were purified using ExoSAP-IT™ PCR Product Cleanup Reagent (Applied Biosystems 78201.1.ML) and sequenced by Sanger (Macrogen). The results of T1 genotyping were used for prioritizing the lines with efficient editing for T2 analysis.

For T2, 2 samples containing 3 seedlings from each of the induction plates each day after confocal imaging were collected into an Eppendorf tube and frozen in liquid nitrogen with two 4mm glass beads. Samples were grinded on a tissue lyser at 30 Hz for 1 min. Tissue powder was diluted in 100 ul of Phire Plant Direct PCR MasterMix dilution buffer (Thermo F160L), 1 ul of the lysate was used for genotyping reaction, using the same primers as for T1 genotyping.

Stable *pat19;20;22* mutant^6^ pheno- and genotyping was performed as described here for the inducible mutant, on 4 days old seedlings. Single nucleotide polymorphism mutations were confirmed with Sanger sequencing (data not shown).

### Chemical treatments

For hydroxylamine (HA) treatment, 7 days old seedlings were transferred for 4 h into 6 well plate filled with liquid ½ MS medium supplemented with 50 mM hydroxylamine (Hydroxylamine hydrochloride, Sigma 159417, 2M stock pH 7.0 in 100 mM HEPES) or 50 mM NaCl (mock), pH 5.8.

For 2-bromopalmitate (2-BP) treatment, 7 days old seedlings were transferred for 15 h (overnight) into 6 well plate filled with liquid ½ MS medium containing 20 µM 2-BP (2-Bromohexadecanoic acid, Sigma 21604) or equivalent volume of DMSO (mock).

After the treatment, seedlings were mounted in a droplet of 20 µg/ml PI stain and imaged under Leica SP8 MP microscope.

### Confocal microscopy of plant lines

Imaging of inducible CRISPR mutants and all the reporters were performed in T2 or later generations, with at least 3 independent T-DNA insertion lines per construct.

Arabidopsis reporter and TurboID lines from Fig. 1-3, 5B-R, S1, S2D, and all Marchantia lines from Fig. 4 were imaged on a Leica SP5 confocal microscope equipped with Apo λ 63×/1.10 water immersion objective, hybrid detectors and an Argon laser. The following laser and hybrid detector settings were used: YFP/mNeonGreen/mCitrine excitation at 514 nm and emission at 520-570 nm; mScarlet/TdTomato excitation at 561 nm and emission at 580-650 nm; mTurquoise excitation at 440 nm (diode) and emission at 460-500 nm. The microscope was controlled by Leica Application Suite: Advanced Fluorescence (LAS AF) software.

Arabidopsis *pat19;20;22* inducible and stable mutants, as well as polar proteins reporters in the background of *pat19;20;22* stable mutant, and reporter lines treated with HA and 2-BP (Fig. 5V-Y, 6, S4, S5), were imaged on a Leica SP8 Multiphoton (MP) microscope equipped with 40x water immersion objective, hybrid detectors and Chameleon MP laser. YFP, mNeonGreen, PI stain and Cas9-RFP were excited with 950 nm 2-photon excitation. Detection was as follows: 510-550 nm (YFP, mNeonGreen), 600-650 nm (PI, RFP). In 2-channel imaging, simultaneous scanning mode was used. The microscope was controlled by Leica Application Suite X (LAS X) software.

### Quantification of PM signal fraction in Arabidopsis roots

Quantification of PM-localized fluorescence signal (Fig. 6 and S5) was performed on 2-channel images obtained with Leica SP8 MP microscope with 40x objective without digital zoom, on full view field. To quantify the fraction of PM-bound protein, images in PI channel were imported to Fiji and processed to create a binary mask using the adaptive threshold by the following Fiji macro command: run(“Auto Local threshold”, “method=Mean radius=15 parameter_1=0 parameter_2=0”). The mask for each image was imported to MATLAB R2021b. The ratio of the total intensity in the PI-rich pixels of the polar protein channel versus the total intensity of the polar protein channel was quantified using the code below. It was referred to as a PM-bound protein fraction.

~~~
C1masked = C1 .* PImask;
C1nonmasked = C1 .* (∼PImask);
PMint = sum(C1masked(:)); % sum in masked region
Totalint = sum(C1(:)); % total polar protein intensity
BoundProtein = 100 * PMint / Totalint;
~~~

Corresponding reporter lines with WT and pat19;20;22 background were all in T2 generation, transgenic plants were selected for seed fluorescent coat when sawed. PM fraction in each independent line was compared using using aov (PM fraction ∼ Genotype) and TukeyHSD functions in R v4.4.2. PM fraction values were visualized using R ggplot2, groups of significant difference were calculated with multcompView R package.

### Human cells – Cell culture

U2OS and HeLa cells were cultured in Dulbecco’s Modified Eagle Medium (DMEM; Capricorn Scientific) supplemented with 10% Fetal Bovine Serum (FBS; Corning) and 100 U/mL Penicillin and 100 µg/mL Streptomycin (1% Pen Strep; Sigma). Cells were kept at 37°C with 5% CO2. Cells were routinely tested for mycoplasma contamination using LT07-518 Mycoalert assay (Lonza).

### Human cells – DNA construct

AtSOK1, AtSOK1[C233S], AtSOK1[C233A], AtPAT19 and AtPAT19[C204S] coding sequences were amplified from *A. thaliana* cDNA (primers listed in Supplementary data 4) and cloned into BamHI site of pCMV:EGFP-N1 or pCMV:mCherry-N1 backbone vectors using NEBuilder® HiFi DNA Assembly Master Mix (NEB E2621S) to create SOK1-EGFP and PAT19-mCherry fusions. DNA for transfection was prepared using Plasmid Midi Kit (Qiagen #12243).

### Human cells – Cell transfection

Cells were plated at 15% confluency on 12 mm cover slips in 12-well plates the day prior to transfection. A ratio of 1 µg plasmid DNA to 3 µL Fugene6 (Promega) transfection reagent was used, transfection mix was prepared in Opti-MEM (Gibco) according to manufacturer’s instructions. Cells were used for fixation and imaging 24 h after transfection.

### Human cells – Imagin

For membrane staining, the DMEM medium in 12-well plates with transfected cells was replaced with DMEM containing 0.5 µg/ml CellMask™ Deep Red Plasma Membrane Stain (Invitrogen C10046). Cells were incubated with the staining medium for 1 min at RT covered in aluminium foil, then the medium was replaced with 4% paraformaldehyde (PFA) in MRB80 buffer (80 mM K-PIPES, pH 6.8, 4 mM MgCl_2_, 1 mM EGTA) with 4% sucrose warmed to 37°C and the cells were fixed for 15 min at RT. After fixation, the wells were washed 3 times with phosphate-buffered saline (PBS) and mounted on glass slides with Mowiol mounting medium (10% Mowiol, Sigma, 81381; 30% glycerol, 60% 0.2 M Tris, pH 8.5).

Fixed cells were imaged on a Confocal Laser Scanning microscope Zeiss LSM 700 using Plan-Apochromat 63x/1.40 Oil DIC (WD=0.19mm) objective. Wavelength settings were the following: EGFP 488 nm laser excitation, LP490 filter emission; mCherry 555 nm excitation, LP560 filter emission; CellMask Deep Red 639 nm excitation, LP640 filter emission. The microscope was used in sequential scanning mode with MA-PMT detector and controlled with ZEN 2011 software.

### Flies

*Drosophila melanogaster* lines were grown on standard cornmeal/agar/molasses media at 25°C. There are no known differences in the physical and molecular mechanisms of planar polarity in male and female flies, thus flies were not distinguished based on sex.

### Generation of transgenic flies

*AtSOK1* was tagged at the C-terminus with mEGFP, and *AtPAT19* was tagged at the C-terminus with 3xHA tags. Both were then cloned into a modified version of the *pActP-FRT-polyA-FRT* vector ^28^ that contains attB recombination sequences. Constructs were integrated into the genome via ΦC31-mediated recombination into the *attP40* or *attP2* landing sites. Transgenics were made by Genetivision. *hs-FLP* was used to excise the *FRT-Stop-FRT* cassette in the germ line.

### Dissection and immunolabeling of prepupal and pupal wing

6 h APF pre-pupal wings were dissected in Schneider’s medium (GIBCO) containing 10% fetal bovine serum. Wings were mounted in a small volume of media containing 1.25% methyl cellulose (Sigma cat# 274429) to reduce sample movement, and imaged within 30 min.

To image live 16 h APF pupal wings, a piece of pupal cuticle above the wing was removed, and pupae were mounted on a glass coverslip in a small drop of halocarbon oil.

For immunolabelling, pupal wings were dissected at 28 h after puparium formation (APF) at 25°C. Briefly, pupae were removed from their pupal case and fixed for 35-40 min in 4% paraformaldehyde in PBS. Wings were then dissected and the outer cuticle removed, and were blocked for 1 h in PBS containing 0.2% Triton X100 (PTX) and 10% normal goat serum. Primary and secondary antibodies were incubated overnight at 4°C in PTX with 10% normal goat serum, and all washes were in PTX. After immunolabeling, wings were post-fixed in 4% paraformaldehyde in PBS for 30 min. Wings were mounted in 25 µl of PBS containing 10% glycerol and 2.5% DABCO, pH7.5 and imaged within 24 h.

### Imagin

Wings were imaged on a Nikon A1R GaAsP confocal microscope using a 60x NA1.4 apochromatic lens, with a pixel size of 80 nm and Z-slices spacing of 200 nm.

### Quantitation of membrane levels and polarity in pupal wing

For measurement of plasma membrane intensity, cytoplasm intensity or membrane to cytoplasm ratio, average projections were generated of ten confocal slices that included the apico-lateral junctional region and seven more lateral slices. Membrane masks for each image were generated in Tissue Analyzer^29^, and custom MATLAB scripts were used to calculate mean membrane intensity, where the mask radius was 3 pixels. Non-masked regions were used to determine mean cytoplasm intensity^30^. Values were normalized to the mean SOK1 membrane intensity in the absence of PAT19.

## Inventory of Supplementary Materials

**Supplementary Figures 1-6**

**Supplementary Table 1**

**Supplementary Data 1-5 (separate files):**

- Supplementary Data 1. Arabidopsis IP-MS sample description and Perseus-processed results, related to Fig. 1.
- Supplementary Data 2. Arabidopsis proximity labeling sample description and Perseus-processed results, related to Fig. 2, 3, 5S and S1.
- Supplementary Data 3. Marchantia proximity labeling sample description and Perseus-processed results, related to Fig. 4.
- Supplementary Data 4. Data underlying the Figures of this study, related to Fig. 4D, 5T-U, 6M, 7K-M, S4B, S5L-N.
- Supplementary Data 5. Transgenic lines, plasmids and primers used in this study.

**Fig. S1.**
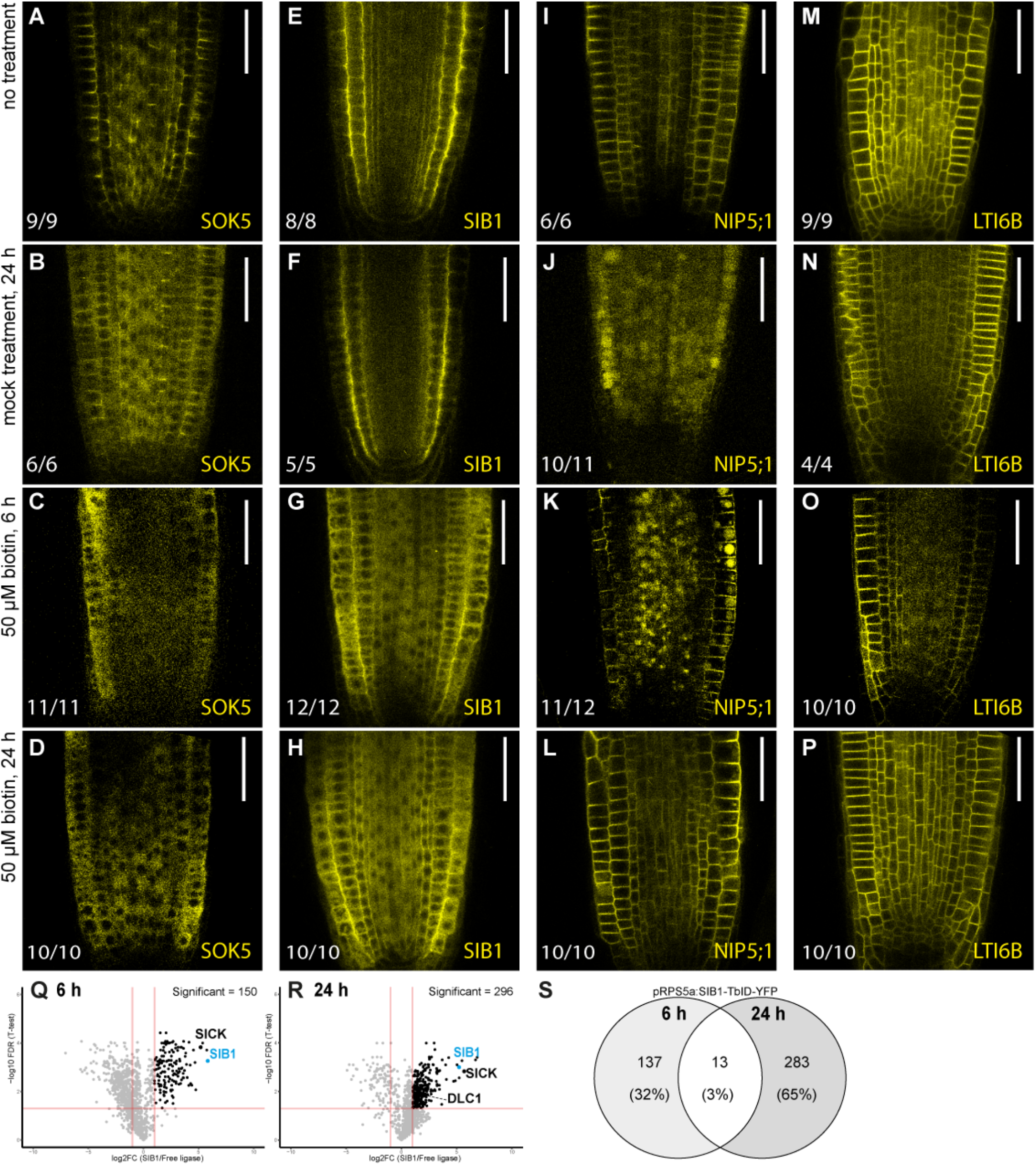
Optimization of biotin treatment for proximity labeling. (**A-P**) Confocal images of 7-day-old roots expressing pRPS5a:SOK5-TbID-YFP (**A-D**), pRPS5a:SIB1-TbID-YFP (**E-H**), pRPS5a:NIP5;1-TbID-YFP (**I-L**), or pRPS5a:LTI6B-TbID-YFP (**M-P**), grown vertically in square Petri plates under standard conditions (**A, E, I, M**), or turned horizontally on day 7 and flooded for 24 h with liquid ½ MS medium (mock; **B, F, J, N**), for 6 h with medium containing 50 µM biotin (**C, G, K, O**), or for 24 h with medium containing 50 µM biotin (**D, H, L, P**). (**Q, R**) Volcano plots showing proteins significantly enriched (Benjamini-Hochberg FDR < 0.05, logFC > 1) in proximity labeling experiments comparing technical triplicates of 7-day-old seedling roots expressing pRPS5a:SIB1-YFP against a free ligase control, treated with 50 µM biotin for 6 h (**Q**) or 24 h (**R**). (**S**) Venn diagram showing the overlap of significantly enriched proteins between (**Q**) and (**R**).

**Fig. S2.**
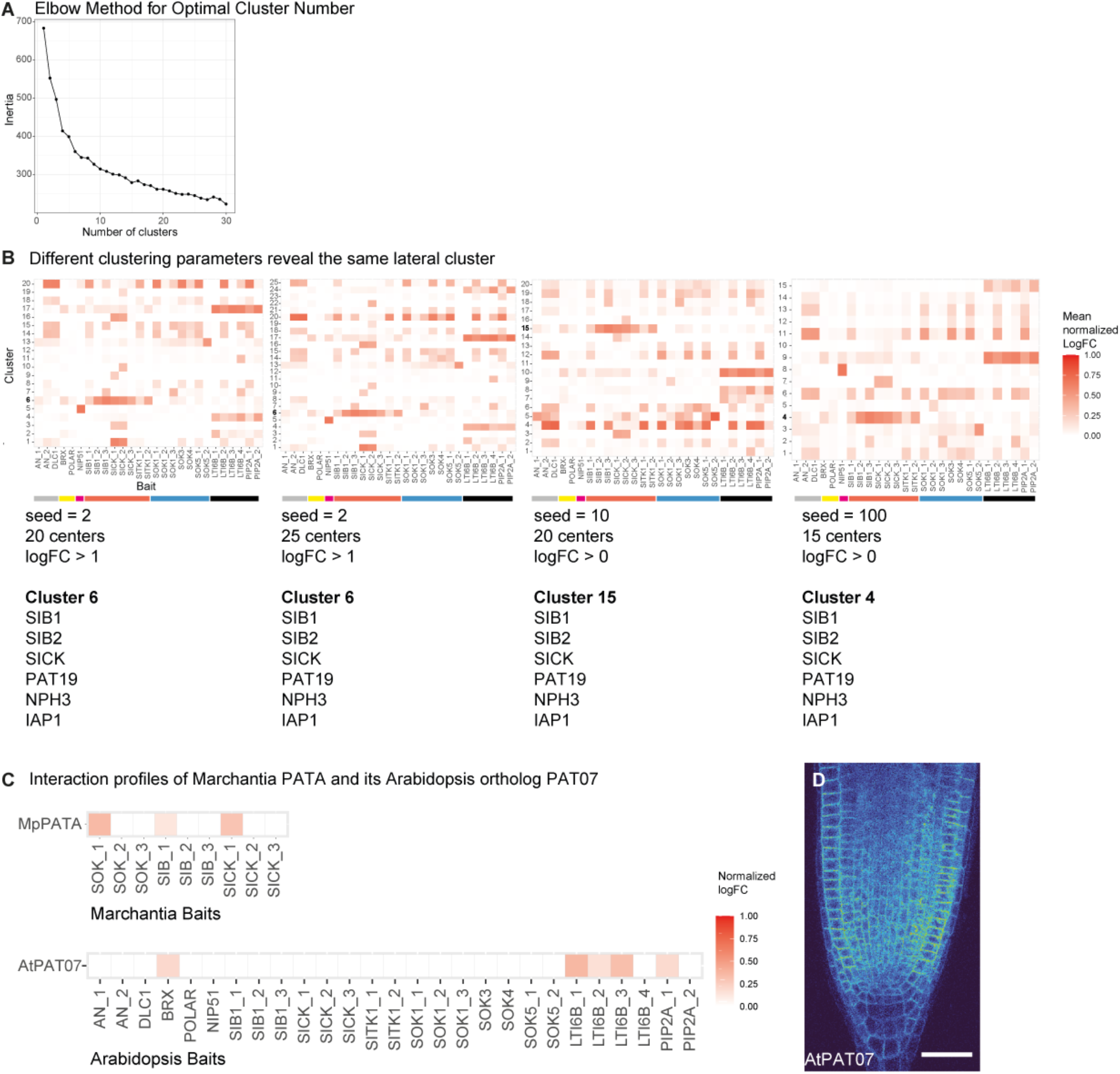
Clustering parameter optimization and cross-species comparison of PAT07 interaction patterns. (**A**) Elbow method used to define the optimal number of k-means clusters (centers) for the interaction profiles of significantly enriched interactors identified in the polar protein proxitome. Inertia reflects how tightly points are grouped around cluster centers; cluster numbers above 15 reach a plateau in inertia and were therefore considered optimal. (**B**) K-means clustering results using different seed parameters and cluster numbers. Bait localization is color-coded as in **Fig. 3**. The heatmap shows mean normalized logFC across all proteins in each cluster; the composition of the laterally enriched cluster remained stable across parameter combinations and is listed below each heatmap. (**C**) Identification of PATA/PAT07 orthologs in the Marchantia and Arabidopsis proximity labeling datasets. Heatmap shows normalized logFC values. (**D**) Confocal image of a 7-day-old root expressing pRPS5a:AtPAT07-TbID-YFP.

**Fig. S3.**
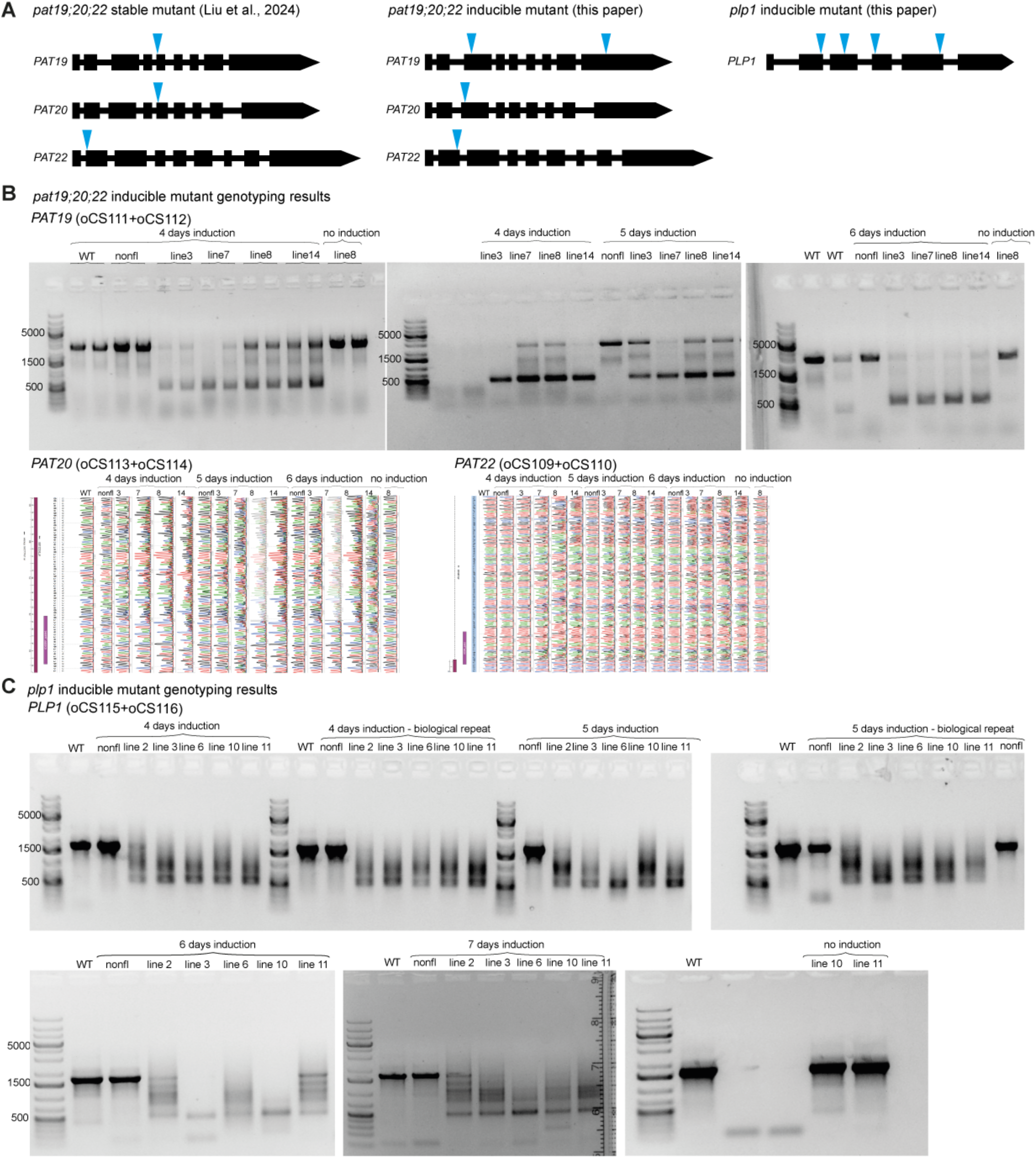
Editing efficiency in inducible CRISPR lines. (**A**) Genomic structure and sgRNA positions (blue arrowheads) for CRISPR mutants used in this study: stable *pat19;20;22*, inducible *pat19;20;22* and *plp1*. (**B**) Genotyping confirms efficient editing in pRPS5a:XVE pLexA:Cas9-RFP *pat19;20;22* after 4-6 days of β-estradiol induction (truncated fragments for PAT19; Sanger sequencing shifts for PAT20 and PAT22). WT – wild type Col-0 control, “nonfl” – non-fluorescent seeds from the same mutant seed batch. (**C**) Genotyping confirms efficient editing in pRPS5a:XVE pLexA:Cas9-RFP *plp1*, shown by truncated PLP1 fragments. **Primers listed in Supplementary data 5**.

**Fig. S4.**
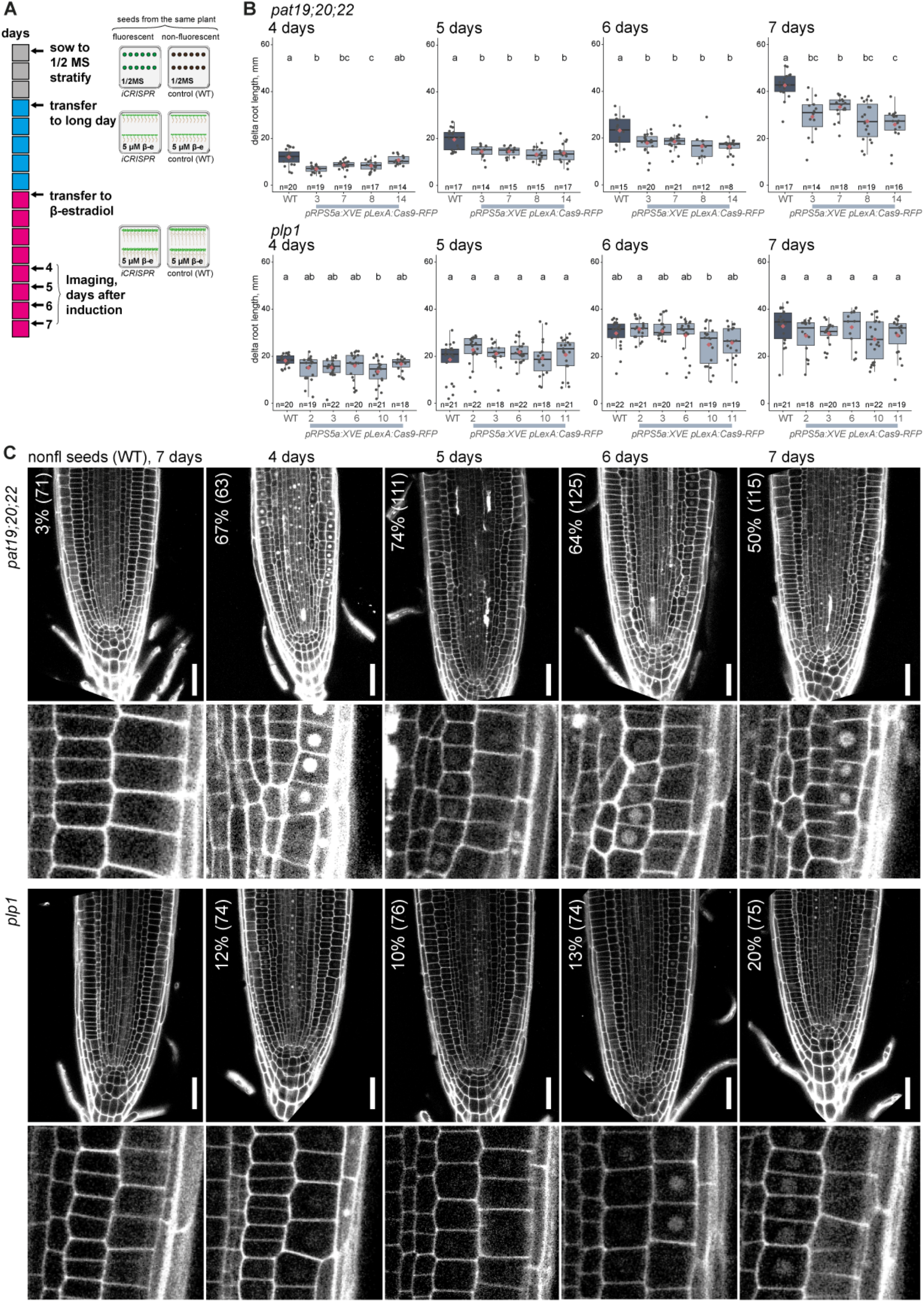
Induction time-course for *pat19;20;22* confirms the involvement of PAT19/20/22 in root growth regulation. (**A**) β-estradiol induction scheme. T2 seeds from independent lines were used for all experiments. (**B**) Boxplots of main root length increase over induction days, comparing non-fluorescent WT seed controls and several independent T2 inducible CRISPR lines for *pat19;20;22* or *plp1*. Boxes show median (black line), 1st/3rd quartile (box borders), and 1.5x interquartile range (whiskers); red diamonds indicate mean; gray dots indicate individual data points. The number of observations per line is shown below each box. Letters denote statistical groups (Tukey HSD post-hoc test). Measurements and p-values are listed in Supplementary data 4. (**C**) Confocal root images of T2 inducible CRISPR plants grown as in (**A**). Roots were stained with propidium iodide; Cas9-RFP was imaged in the same channel. Numbers indicate the percentage of roots with cell division defects and the total number of roots analyzed. Complete data, including the numbers of roots exhibiting cell division defects per line, are provided in **Supplementary Data 4**. Insets below each image show normal (WT, *plp1*) or disturbed (*pat19;20;22*) cell division orientation. Scale bar: 40 µm.

**Fig. S5.**
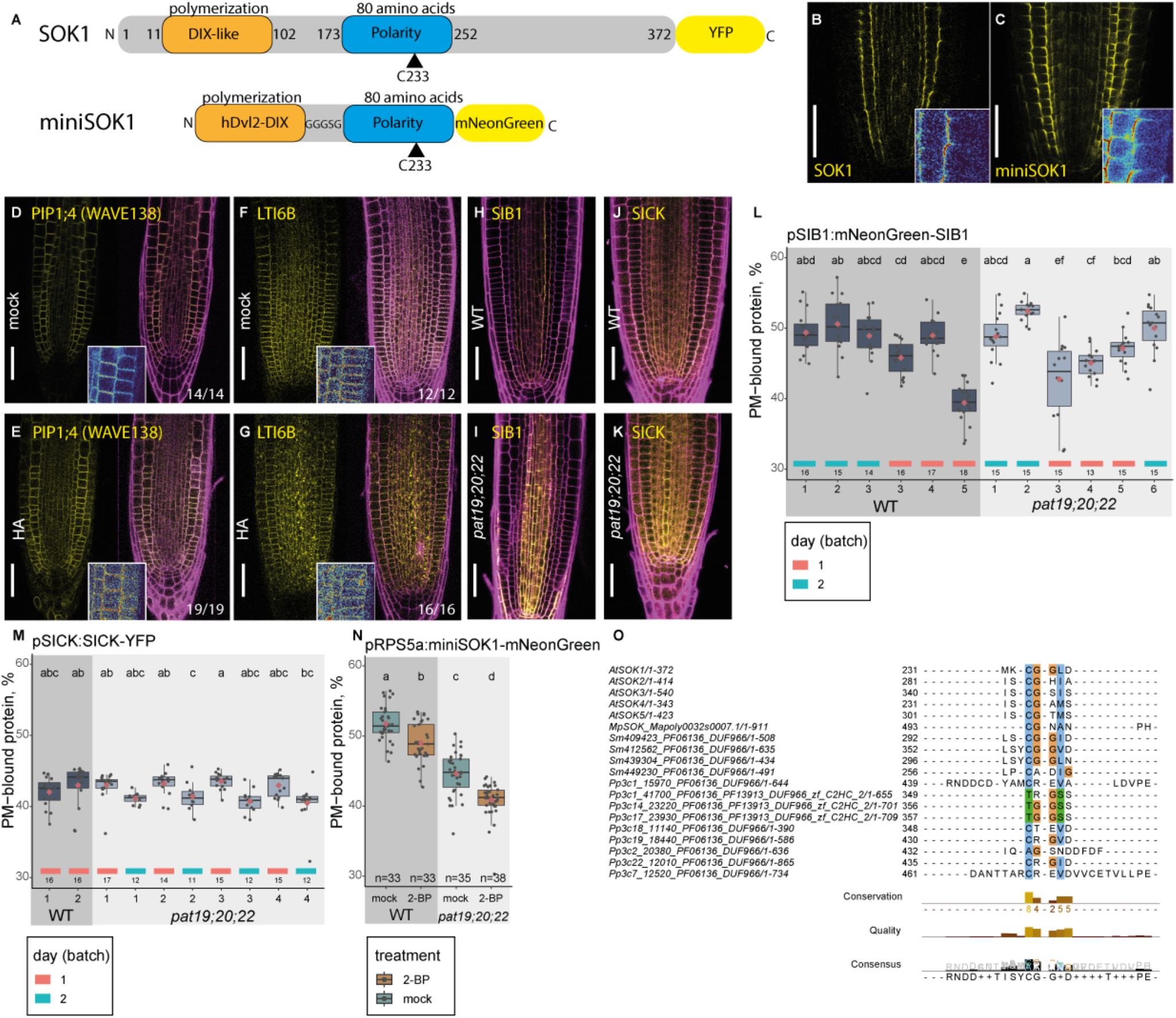
PAT19 is required for SOK1, but not for SIB1 or SICK, PM association. (**A**) Domain structure of full-length Arabidopsis SOK1 and the minimal polarity chimeric reporter, miniSOK1. (**B-C**) Confocal images of 7-day-old roots expressing pRPS5a:SOK1(full-length)-YFP (**B**) or pRPS5a:miniSOK1-mNeonGreen (**C**). Bottom-right inset in each image shows a few epidermal and cortex cells from the right side of the root, false-colored by intensity. (**D-G**) Confocal images of 7-day-old roots expressing pUB10:PIP1;4-EYFP (WAVE131; **D, E**) or pRPS5a:LTI6B-TbID-YFP (**F, G**), treated for 4 h with 50 µM hydroxylamine (HA; **E, G**) or 50 µM NaCl (mock; **D, F**). Each panel shows the protein of interest (yellow) and a merge with propidium iodide (PI) cell wall stain (magenta); bottom-right inset shows false-colored intensities. (**H-K**) Confocal images of 7-day-old roots expressing pSIB1:mNeonGreen-SIB1 (**H, I**) or pSICK:SICK-YFP (**J, K**) in WT (**H, J**) or stable *pat19;20;22* mutant (**I, K**) backgrounds. (**L-M**) Quantification of the PM-associated fraction of fluorescent signal for SIB1 (**L**) and SICK (**M**) lines shown in (**H-K**). Boxes show median (black line), 1st/3rd quartile (box borders), and 1.5x interquartile range (whiskers); red diamonds indicate mean; gray dots indicate individual data points. The number of observations per line is shown below each box. Letters denote statistical groups (Tukey HSD post-hoc test). Imaging was performed on different experimental days, indicated by colored tiles below the boxes. Raw data and p-values are in **Supplementary data 4**. (**N**) Quantification of the PM-associated fraction of fluorescent signal for miniSOK1 lines under 2-BP treatment, shown in **Fig. 6N-Q**. Boxplot statistics and data availability are as in (**L, M**). Scale bars: 40 µm. (**O**) A fragment of multiple sequence alignment of plant SOSEKI sequences showing a conservation of AtSOK1[C233] with MpSOK[C493]. Sequences of *Arabidopsis thaliana* (At), *Marchantia polymorpha* (Mp), *Selaginella mollendorfii* (Sm) and *Physcomitrium patens* (Pp) homologs.

**Fig. S6.**
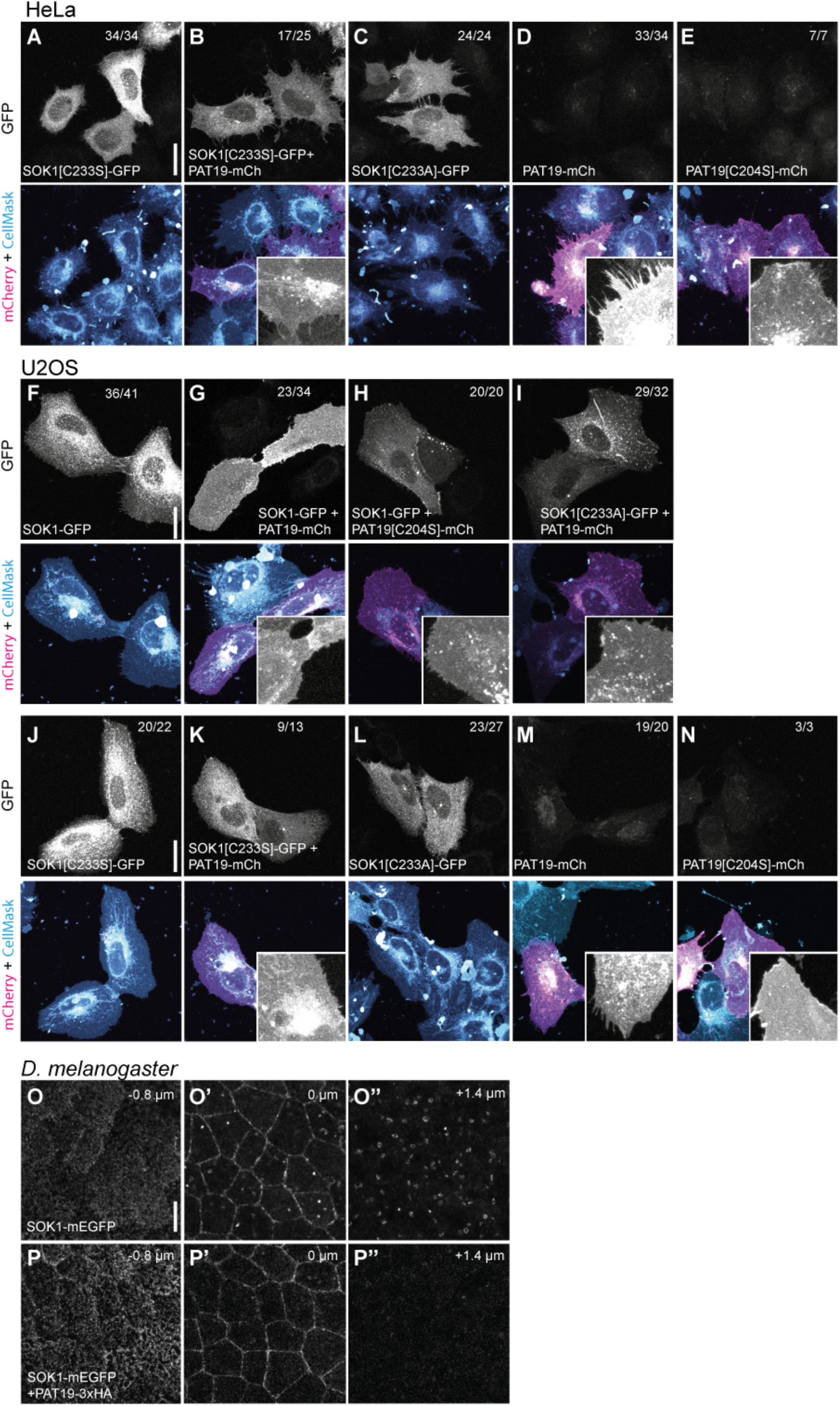
PAT19 alters subcellular SOK1 localization in heterologous human cell systems. (**A-E**) Maximal intensity projections of confocal images of fixed HeLa cells transiently expressing combinations of full-length SOK1-GFP, PAT19-mCherry, and their cysteine mutants (SOK1[C233S/A], PAT19[C204S]). Membranes were stained with CellMask Deep Red. Insets show a fragment of a cell with PAT19-mCherry. Numbers indicate the fraction of cells showing the SOK1/PAT19 localization pattern in the representative image, across 2 independent experiments. Scale bar: 20 µm. (**F-N**) As in (**A-E**), but in U2OS cells. (**O-P**) PAT19 promotes association of SOK1-mEGFP to apical and apico-lateral regions of Drosophila epithelia. mEGFP fluorescence in 16 hr APF live prepupal wings expressing SOK1-mEGFP (**O**) or SOK1-mEGFP and PAT19-3xHA (**P**). Confocal sections covering the apical cell surface. (**O’**,**P’**) Confocal sections that are 0.8 µm below, at the level of the apico-lateral cell junctions. (**O’’**,**P’’**) Confocal sections that are a further 1.4 µm more basal. Scale bar: 5 µm.

**Table S1.**
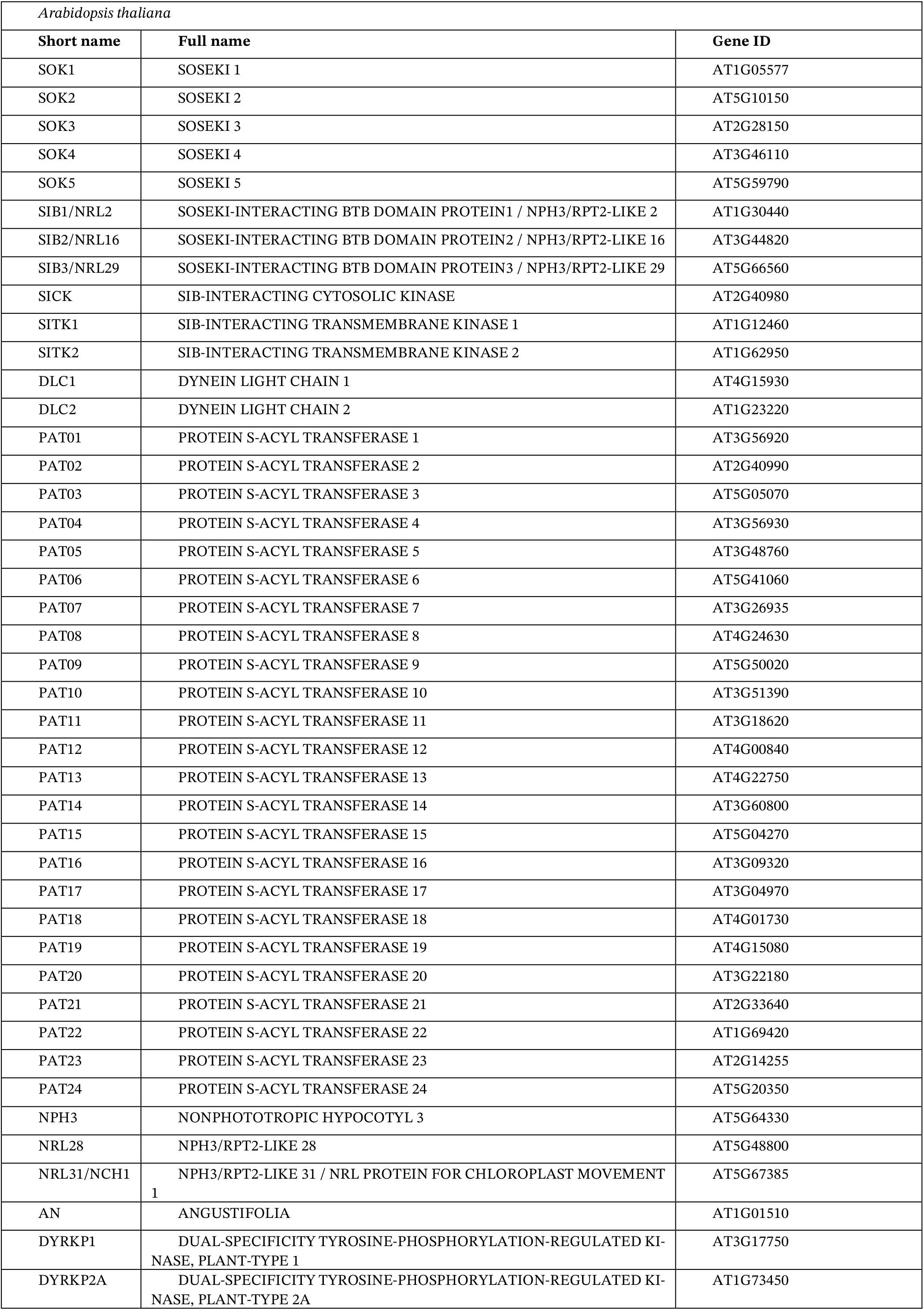
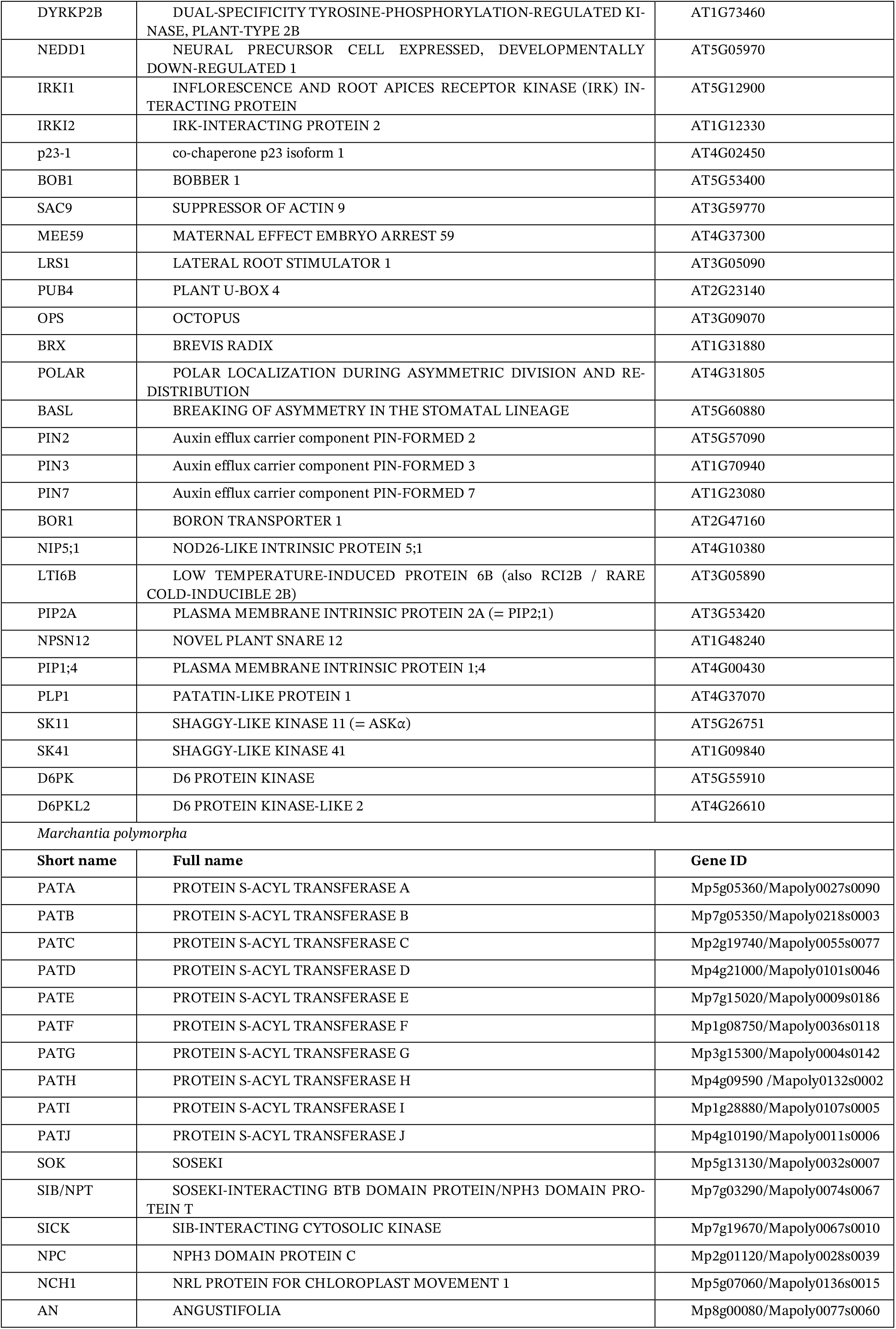
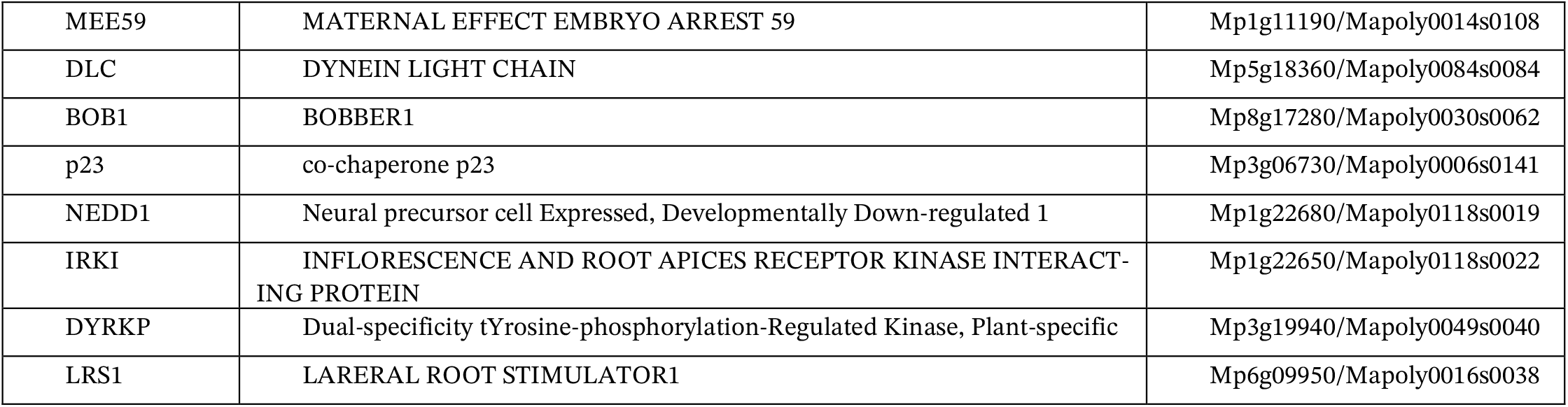
Gene IDs and trivial protein names used in this study.

